# *SpaceTrooper*: a quality control framework for imaging-based spatial omics data

**DOI:** 10.64898/2025.12.24.696336

**Authors:** Benedetta Banzi, Dario Righelli, Matteo Marchionni, Oriana Romano, Mattia Forcato, Davide Risso, Silvio Bicciato

## Abstract

Quality control (QC) is a critical step in the analysis of imaging-based single-cell spatial omics data, yet standardized metrics tailored to these technologies are still lacking. Most existing QC approaches are adapted from single-cell sequencing workflows and rely on fixed thresholds, limiting their ability to capture complex artifacts arising from image processing, tissue morphology, and platform-specific effects. Here, we present SpaceTrooper, a data-driven QC framework that computes an integrated per-cell quality score by combining expression-derived and morphological features. Without relying on fixed thresholds, SpaceTrooper systematically identifies low-quality cells caused by segmentation errors, signal loss, elevated background, spatial distortions, and tissue-derived artifacts. Across diverse tissues, technologies, and modalities, SpaceTrooper robustly detects technical failures and substantially improves clustering and cell-type coherence relative to conventional QC strategies.

## INTRODUCTION

Single-cell spatial omics technologies enable the quantitative profiling of transcripts and proteins directly within intact tissues, preserving the native spatial organization of cells while capturing high-dimensional molecular information^1–6^. By jointly resolving molecular states and spatial context at single cell resolution, these approaches provide unprecedented insight into cellular identities, tissue architecture, and multicellular organization within complex biological systems^7–9^. Integrating molecular, cellular, and spatial information further reveals how local interactions, spatial gradients, and tissue structure shape gene expression programs and cellular behavior, offering a powerful framework to study development, homeostasis, and disease^8,10–14^.

The rapid expansion of single-cell spatial omics has generated datasets of increasing size, complexity, and heterogeneity, creating analytical challenges that extend beyond the scope of computational workflows developed for dissociated single-cell data. This has motivated the development of dedicated methods that integrate molecular profiles with spatial coordinates to infer cell identities, delineate tissue organization, and characterize spatially structured cell-cell interactions^15–24^. However, while downstream analytical methods have advanced rapidly, quality control (QC) strategies for imaging-based spatial omics remain comparatively underdeveloped and lack standardization^25–31^.

Imaging-based spatial omics relies on fluorescence in situ hybridization-based detection of molecular targets and requires accurate cell segmentation to assign signals to individual cells and generate spatially resolved expression matrices^32^. Errors arising from tissue damage, segmentation inaccuracies, signal loss, elevated background, or image acquisition artifacts can introduce low-quality cells that propagate technical noise into downstream analyses. Current approaches to identify such cells typically rely on visual inspection or fixed thresholds adapted from single-cell sequencing workflows and applied to technology- or modality-specific metrics^25–31^. Visual inspection is subjective and difficult to scale, whereas threshold-based filtering applies individual metrics independently across cells, overlooking cell-to-cell variability and potentially confounding technical artifacts with genuine biological variation. As spatial atlases continue to expand across tissues and molecular modalities^33^, there is a growing need for quantitative, reproducible, and scalable QC frameworks that operate at single-cell resolution and integrate complementary sources of quality information.

Here we present SpaceTrooper, a technology-independent quality control framework for imaging-based spatial omics data. SpaceTrooper integrates morphological descriptors and expression-derived features into a data-driven model that computes a per-cell quality score (QS), enabling systematic identification of low-quality cells without reliance on predefined thresholds. The QS captures diverse sources of technical failure, including segmentation errors, tissue-altered regions, weak molecular signal, elevated background noise, and spatial distortions related to image acquisition. We demonstrate the applicability of SpaceTrooper across RNA- and protein-based imaging spatial omics datasets generated using CosMx Spatial Molecular Imaging (NanoString), Xenium (10x Genomics), and MERFISH (Vizgen), spanning diverse tissues and experimental conditions, and show that filtering based on the integrated quality score improves downstream analyses compared with individual QC metrics. SpaceTrooper is released as an open-source R/Bioconductor package built on the SpatialExperiment infrastructure^34^, enabling seamless integration into existing spatial omics analysis workflows.

## RESULTS

### Overview of SpaceTrooper QC framework

The SpaceTrooper QC framework quantifies a composite cell quality score (QS) by integrating multiple complementary metrics that capture distinct technical artifacts in imaging-based single-cell spatial omics data (**Figure 1A**). Input data, including gene expression matrix, cell-level metadata, cell boundary polygons, and, when available, field-of-view (FOV) coordinates, are standardized into a unified *SpatialExperiment* (SPE) object, providing a robust, platform-agnostic foundation for data harmonization, quality assessment, and downstream analyses^34^. We defined the cell QS using a set of complementary cell-level metadata that are consistently available across imaging-based spatial omics datasets generated with the CosMx (NanoString), Xenium (10x Genomics), and MERFISH (Vizgen) platforms (**Supplementary Table 1**). These datasets span RNA and protein assays at multiple panel coverages in both normal and pathological tissues from different species. The selected metrics capture distinct yet interrelated aspects of data quality, including morphology, molecular content, background noise, and spatial distortion (**Figure 1A** and **Supplementary Table 1**; see **Supplementary Note 1** for details on the selection of the quality score components). Cell size (area or volume) reflects cell morphology and segmentation quality, and its distributions in the analyzed datasets exhibit a right-skewed shape with a pronounced tail toward extreme values (**Supplementary Figures 1A, 1C, 1E, and 4A**). Within the biologically plausible size range, cell size is approximately linearly associated with total molecular counts, whereas this relationship is not preserved at extreme values, consistent with segmentation artifacts (**Supplementary Figures 1B, 1D, 1F, and 4B**). Signal density, defined as molecular counts normalized by cell size, displays heterogeneity across samples and enables the selective identification of low-density outliers in the left tail of the distribution while preserving genuinely small cells (**Supplementary Figures 2A-F and 4C**). Background signal, quantified as the proportion of control probe counts, is generally low but right-skewed across all platforms, and cells with elevated background fractions tend to be associated with low signal content (**Supplementary Figures 3A-F and 4D-E**). For NanoString CosMx, but not for Xenium or MERFISH datasets, we observed systematic FOV border-related artifacts. Both cell area and total probe counts decrease as cells approach the nearest FOV border (**Supplementary Figures 5A-B and 6A-B**). To capture these distortions, we leveraged the cell aspect ratio as a geometric proxy, which remains close to unity for inner cells but deviates sharply near horizontal and vertical borders due to truncated cell height or width (**Supplementary Figures 5C-D and 6C-D**).

**Figure 1.**
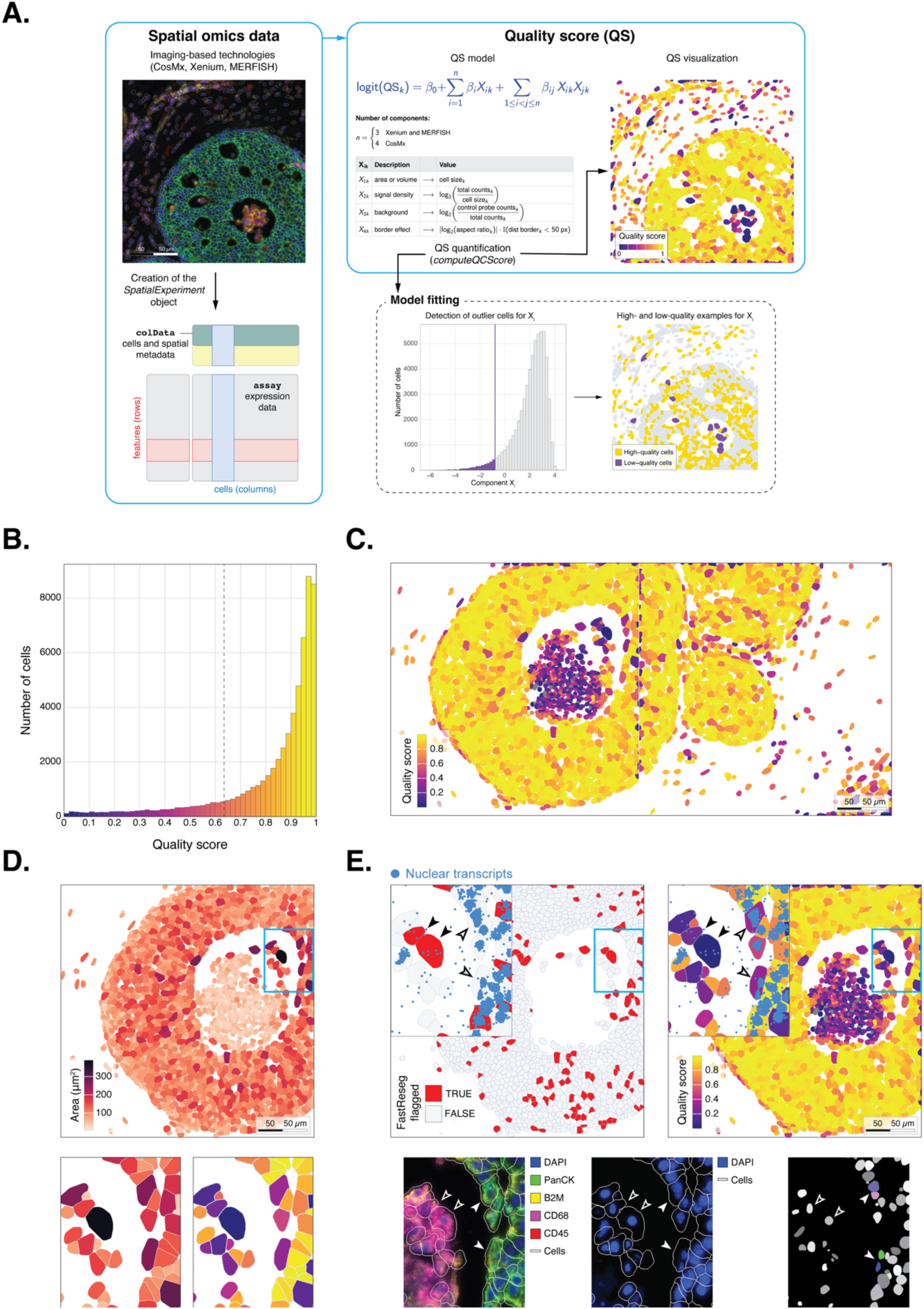
SpaceTrooper detects low-quality cells in imaging-based spatial omics data. **A.** Overview of the SpaceTrooper QC framework. Imaging-based spatial omics data are stored in a *SpatialExperiment* object containing cell-level metadata and the assay expression matrix. For each cell *k*, the cell quality score (QS) is computed as the log-odds of being high- versus low-quality using a ridge-regularized logistic regression model that integrates *n* predictor components, i.e., cell size, signal density, background signal, and, for CosMx, border-effect distortion. High- and low-quality examples are defined by adaptive, distribution-based outlier detection, and used to fit the model. The resulting QS is a continuous probability (ranging from 0 to 1) that reflects cell-level data quality. **B.** Distribution of QS values in the CosMx 1k breast cancer dataset, showing a left-skewed profile with a distinct tail of low-quality cells. The vertical dashed line indicates the median minus 3 MAD threshold. **C.** Spatial mapping of QS across FOV 11 and 12, highlighting the enrichment of low-quality cells within ductal lumen and along FOV boundaries. **D.** Spatial distribution of cell area (in μm^2^) in FOV 11 (top). The inset zooms show enlarged polygons at the lumen-epithelium interface and their corresponding QS values (bottom). **E.** Spatial mapping of spatial multiplets detected by FastReseg and of QS (top). In the insets, blue dots mark MALAT1 and NEAT1 nuclear transcripts. In the bottom zooms, composite fluorescence image (left), DAPI fluorescence (middle), and StarDist-labeled nuclei (right). Black-filled arrowheads indicate low-quality cells flagged by FastReseg as spatial multiplets; white-filled arrowheads mark low-QS cells not detected by FastReseg that contain multiple nuclei, consistent with same-cell-type doublets. In the QS plots (panels **C** and **E**), cells shown in grey correspond to cells with zero counts.

For each cell *k*, we compute QS as the log-odds (logit) of being high- versus low-quality based on *n* predictor components:

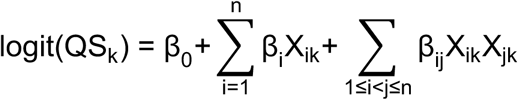

Here, *X_1k_* corresponds to cell area for CosMx and Xenium and to cell volume for MERFISH; *X_2k_* represents the signal density, defined as total probe counts normalized by cell size (and log_2_-transformed) for RNA platforms and as the log_2_-transformed sum of average protein intensities for protein assays; *X_3k_* quantifies the background signal as the log_2_-transformed proportion of control probe counts; and *X_4k_*, included only for NanoString CosMx, captures border-induced distortions through the log_2_-transformed aspect ratio for cells located close to the nearest FOV boundary (e.g., within 50 pixels; **Figure 1A**). For each dataset, the cell quality score is quantified by fitting a penalized generalized linear model (GLM) with a binomial distribution and logistic link, modeling the probability of each cell being high-quality as a function of the selected predictor components. Model coefficients are learned using ridge regularization to mitigate overfitting while retaining all predictor components, with the optimal regularization parameter selected by cross-validation. The example set for estimating the model parameters is constructed defining low- and high-quality cells independently for each component using adaptive, distribution-based outlier detection as Medcouple^35,36^ or Median Absolute Deviation (MAD), depending on distribution skewness (**Supplementary Table 2** and **Supplementary Figures 1-6**). Low-quality examples are those flagged as outliers for at least one component metric, whereas high-quality examples are selected from the central or upper quantiles (depending on the specific component’s characteristics) and, for CosMx, from interior FOV regions (**Figure 1A**; see ‘Quantification of the cell quality score’ in Methods for more details). Components associated with fewer than 0.1% low-quality examples are excluded from model fitting. The resulting QS corresponds to the estimated probability of a cell being high quality (**Figure 1B** and **Supplementary Figure 7**).

### SpaceTrooper detects diverse segmentation, tissue-derived, and imaging artifacts in spatial omics datasets

To demonstrate its ability to identify artifacts in spatial transcriptomics data, we applied the SpaceTrooper QC framework to a dataset obtained from a single tissue section of human ductal carcinoma in situ (DCIS) profiled using the NanoString CosMx Spatial Molecular Imaging platform (CosMx 1k breast cancer dataset). The distribution of the quality score (QS) indicates that most cells exhibit high technical quality (**Figure 1B**). Spatial mapping of the quality score across the DCIS tissue section reveals a heterogeneous distribution of low- and high-quality cells (**Supplementary Figure 8**). High-quality cells predominantly localize within compact epithelial structures whereas low-quality cells cluster within the luminal regions of ductal structures (e.g., FOVs 11, 13, and 22-23), in localized foci, and along tissue or FOV boundaries. A closer inspection of the ductal structure in FOV 11 and the contiguous FOV 12 illustrates how the QS captures differences between biologically meaningful organization and localized technical artifacts. The highest QS values are concentrated in the well-structured epithelial layers, whereas low-QS cells predominate within the lumen and at the borders of the FOVs, consistent with a border effect (**Figure 1C**). Within the lumen, low-QS cells arise from different combinations of component-level deficiencies: some display inflated area values despite average signal density, suggesting oversized segmented objects (**Supplementary Figure 9A**), while others exhibit uniformly reduced molecular content, characterized by low signal density and elevated background (**Supplementary Figures 9B-C**). The aspect ratio component further highlights morphological distortions at the FOV boundary (**Supplementary Figure 9D**), where cell morphologies become fragmented or elongated, reflected in locally low QS values.

Several cells with large area values appear in the upper-right region of FOV 11 (**Figure 1D**), at the interface between the central lumen and the surrounding epithelial mass. As expected, these oversized cells receive low QS values, reflecting their outlying morphological profiles. Because unusually large polygons often arise from the erroneous segmentation of multiple cells, we used FastReseg^37^, which detects cells with divergent transcriptional profiles, to assess whether a subset of low-QS cells corresponds to spatial multiplets. The QS successfully highlighted several large polygons that FastReseg classified as spatial multiplets (black-filled arrows in **Figure 1E**). However, additional large low-QS cells were not flagged by FastReseg, consistent with their interpretation as homo-doublets, i.e., two cells of the same type incorrectly segmented as a single object (cancer cells indicated by white-filled arrows in **Figure 1E**). This explanation is supported by the presence of uniform, high keratin signal (green fluorescence) and multiple nuclei (blue fluorescence and segmented nuclei) in these polygons, features indicative of same-cell-type doublets that are readily detected by the QS. SpaceTrooper also captures artifacts induced by the sample histopathological characteristics. Examination of the ductal lumen revealed a central region consistent with comedonecrosis, a common feature in DCIS breast tumors^38–40^, associated to dense accumulations of pyknotic (i.e., small, dense, and hyperchromatic) DAPI-positive debris, an absence of cytokeratin signal, and a near-complete loss of transcript (**Supplementary Figure 10A**). This necrotic core shows a breakdown of cellular integrity and is bordered by focal clusters of CD68-positive cells, consistent with the presence of macrophages clearing cellular remnants^41^. Within this region, segmented cells contain almost no detected transcript probes, reflected by their markedly reduced RNA signal density and transcript spatial distribution. The low signal density is accompanied by elevated background values, indicating increased noise and degraded signal content. Accordingly, these cells receive very low QS values (**Figure 1C**).

In addition to necrotic regions, loss of RNA signal can also arise from purely technical artifacts, as out-of-focus imaging caused by partial tissue detachment during the multiple hybridization and wash cycles. A portion of FOV 31 displays diffuse, blurred fluorescence signal in the composite and DAPI channels, indicating a localized failure in image focusing (**Supplementary Figure 10B**). Although many segmentation tools fail when nuclei and membranes lose sharp boundaries, some algorithms may still generate polygons in these regions, resulting in segmented cells with severely reduced transcriptional content^42^. Accordingly, cells lying within the out-of-focus area exhibit markedly low signal density and are assigned consistently low QS values (white-filled arrows in **Supplementary Figure 10B**). Furthermore, out-of-focus regions can be associated with local tissue mechanical stress and give rise to cells that, independently of their phenotype, display distorted nuclei, irregular or spindled cell shapes, and reduced intensity in non-nuclear morphological markers. These abnormalities are clearly visible in cells surrounding the blurred zone (grey-filled arrows in **Supplementary Figure 10B**), which exhibit atypical morphologies and depleted RNA signal. Consistently, these cells receive low QS values.

Finally, SpaceTrooper highlights FOV border-related distortions, including artifacts introduced when cells are partially truncated by the boundaries of the acquisition system (**Supplementary Figure 11**). In some cases, cells overlapping FOV borders are partially duplicated across adjacent fields, as illustrated by the two cell sets with high aspect ratios and low QS highlighted at the boundary between FOVs 38 and 39. In the composite immunofluorescence images, the duplicated polygons exhibit similar shapes and identical staining patterns; nevertheless, they are assigned discordant phenotype labels by InSituType annotation. For example, the CD68-positive myeloid cell on the left appears on both sides of the boundary, yet one copy is classified as a mixed breast cancer/TAM while the other is labeled as a mural cell. All together these examples show that the QS provides a unified QC layer that pinpoints diverse sources of artifact that would otherwise confound spatial omics data interpretation.

### SpaceTrooper identifies low-quality cells impacting downstream analysis

Accurate identification and exclusion of low-quality cells is a critical prerequisite for ensuring the reliability of downstream analyses in single-cell studies^43^. This step is particularly important in imaging-based spatial omics, where biological signals are often obscured by high optical noise, sparse transcript detection, and segmentation artifacts^2,44,45^. To evaluate how filtering cells based on the quality score affects downstream results, we computed clusters and UMAP embeddings for the CosMx 1k breast cancer dataset, comparing analyses performed on the full dataset (**Figures 1B** and **Supplementary Figure 8**) and on subsets obtained excluding low-quality cells.

In the analysis of the full dataset, mapping QS values onto the UMAP embedding revealed a distinct aggregation of cells with uniformly low scores (**Figure 2A**). Notably, these cells grouped into the same cluster (**Figure 2B**; cluster 12) whose QS values were significantly lower than those of any other cluster (**Figure 2C**). Because the QS is similarly distributed across cell types (**Supplementary Figure 12A**), a cluster with markedly lower scores is likely indicative of technical artifacts rather than of a biologically meaningful phenotype. To test this, we inspected the distribution of InSituType cell types within the UMAP embedding, revealing a mixture of phenotypes in the same region of cluster 12 (**Figure 2D**). Additionally, these cells exhibited reduced assignment probabilities from InSituType, with cluster 12 showing significantly lower cell type probability compared to any other cluster (**Supplementary Figures 12B-C**). The composition of cluster 12 is highly heterogeneous, including substantial proportions of immune, endothelial, stromal, and epithelial cells. This promiscuity, both in terms of lineages and cell types (**Figure 2E** and **Supplementary Figure 12D**), further supports the notion that cluster 12 is driven by technical noise rather than biological signal. To more rigorously assess the potential impact of this intermingling on downstream analyses, we computed the homophily score, a per cell metric that is independent of UMAP embedding and clustering. The homophily score measures cell-type coherence within the local neighborhood of each node in the k-nearest neighbors (kNN) graph underlying cluster computation, by measuring the proportion of a cell’s nearest neighbors that share the same cell-type label. As expected, the mixed region observed in the UMAP embedding (**Figure 2D**) was characterized by low homophily scores (**Figure 2F**), with cluster 12 exhibiting significantly lower homophily than any other cluster (**Figure 2G**). Together, these results indicate that the QS flags cells that aggregate into artifactually mixed clusters and biologically non-specific regions in UMAP embeddings. Importantly, the presence of these low-quality cells has a global impact on downstream analyses, as they distort the underlying kNN graph structures, thereby affecting both dimensionality reduction and clustering.

**Figure 2.**
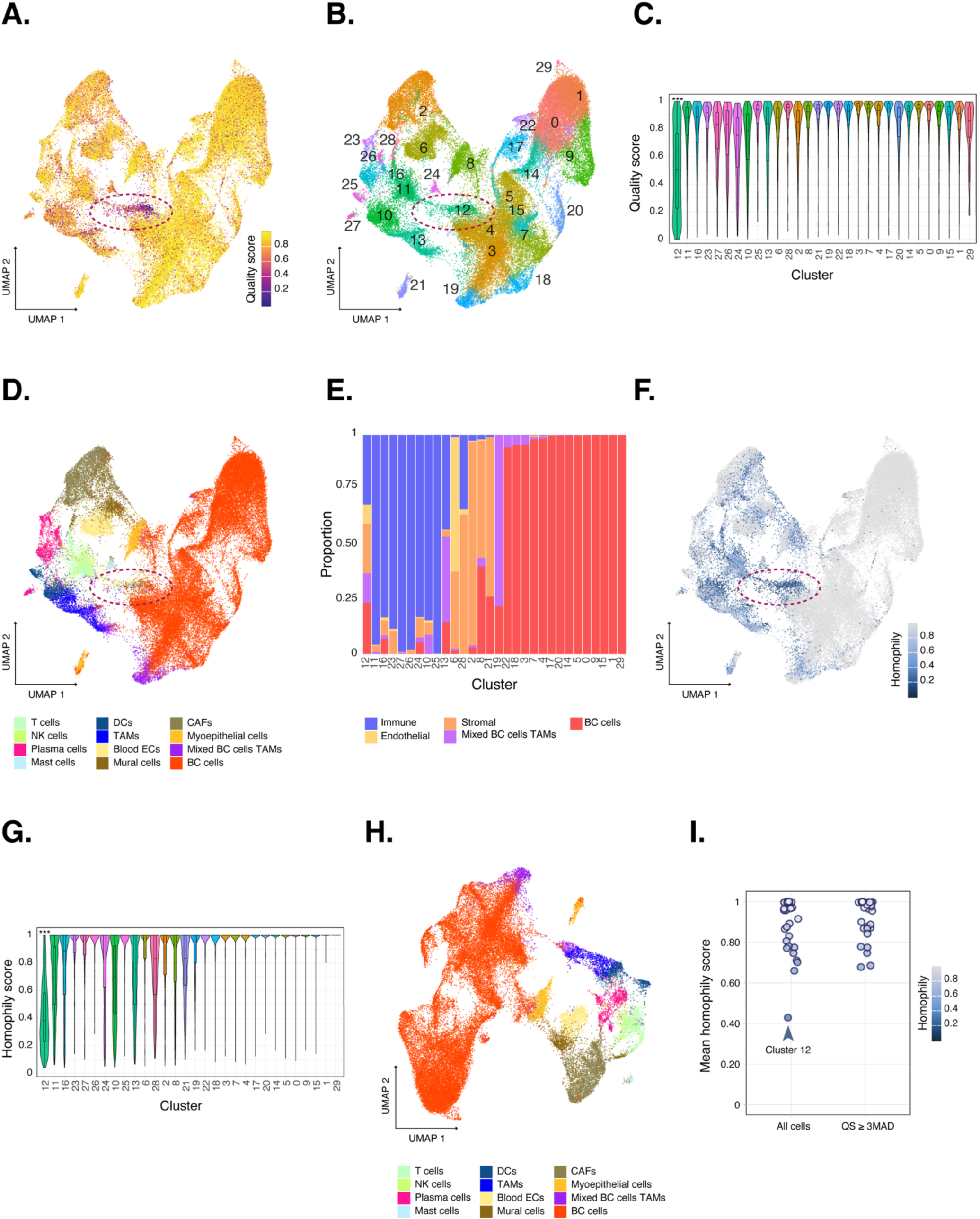
SpaceTrooper identifies low-quality cells impacting downstream analysis. **A.** UMAP embedding of the complete CosMx 1k breast cancer dataset colored by SpaceTrooper quality score (QS). A distinct aggregation of cells with low QS is observed in the center of the embedding (dashed ellipse). **B.** Same UMAP embedding of panel **A** colored by unsupervised clustering, with cluster identities indicated. The low-QS region corresponds predominantly to cluster 12 (dashed ellipse). **C.** Distribution of QS values across clusters. Clusters are ordered according to cell type composition as described in the Method section. Cluster 12 shows significantly lower QS values compared with each of the other clusters (one-sided Wilcoxon test with Benjamini–Hochberg correction; *** p<0.001 for all comparisons). **D.** Same UMAP embedding of panel **A** colored by cell type annotation. The region corresponding to cluster 12 contains a heterogeneous mixture of immune, endothelial, stromal, and breast cancer cell types. **E.** Stacked bar plot showing the proportion of major cell lineages per cluster. Cluster 12 displays marked lineage promiscuity. Clusters are ordered as in **C**. **F.** Same UMAP embedding of panel **A** colored by homophily score, which quantifies cell-type coherence within local kNN graph neighborhoods. The mixed central region exhibits low homophily (dashed ellipse). **G.** Distribution of homophily scores across clusters. Cluster 12 shows significantly lower homophily values compared with each of the other clusters (one-sided Wilcoxon test with Benjamini–Hochberg correction; *** p<0.001 for all comparisons). Clusters are ordered as in **C**. **H.** UMAP embedding after filtering low-quality cells using a data-driven QS threshold based on 3MAD outlier detection method. Removal of low-QS cells results in clearer separation of cell types and elimination of artifactual mixed regions. **I.** Mean homophily score per cluster before and after QS-based filtering. Removal of low-QS cells eliminates the cluster characterized by low homophily.

Building on these observations, we removed low-quality cells using a data-driven QS threshold before downstream analyses. Specifically, we filtered out 8,404 cells (14.18% of the total) with QS values below the median minus 3 MAD (QS threshold = 0.635; median = 0.9154; MAD = 0.094; **Figure 1B**). Visual inspection of the resulting UMAP embedding revealed a clearer separation of cell types, particularly among non-malignant cells, with no evident mixed regions (**Figure 2H**). Consistently, cluster composition showed improved coherence of both cell lineages and cell types (**Supplementary Figures 12E-G**), with no cluster displaying pronounced heterogeneity. To quantify cell label consistency, we computed homophily scores within the kNN graph. Although some cells with low homophily remained, no cluster was dominated by these cells (**Supplementary Figures 12H-I**). Comparing these homophily scores with those previously computed on the full, unfiltered dataset further highlights the benefit of QS-based filtering for improving the reliability of downstream analyses (**Figure 2I**).

### Comparison with fixed-threshold QC approaches

Building on the previous analysis, we compared, in the CosMx 1k breast dataset, QS-based filtering with a more conventional approach, following the AtoMx pipeline recommended by NanoString (**Supplementary Note 1**). We selected three exclusion criteria to identify low-quality cells: i) fewer than 20 detected probes (*total probe counts* filter); ii) a proportion of control probes on total probes exceeding 0.01 (*control probe ratio* filter); and iii) a cell area above 216.14 µm^2^, determined using the median plus 3 MAD to avoid arbitrary thresholds (*area* filter). Each criterion was applied independently and in combination (*combined* filtering criterion). In total, 1,345 cells were filtered based on *total probe counts*, 709 based on the *control probe ratio*, 1,189 based on cell *area*, and 3,185 using the *combined* filtering criterion. The impact of each filter varied across cell types (**Supplementary Figure 13A**); while the *area* filter affected cell types similarly, the remaining filters disproportionately excluded non-malignant cells. Notably, the overlap between cells removed by different individual criteria was minimal, with an intersection over union (IoU) of 0.025 between *total probe counts* and *control probe ratio* and of 0.004 between *control probe ratio* and *area* (**Supplementary Figure 13B**). The low overlap between *total probe counts* and *control probe ratio* likely reflects dropout effects in control probes, which were detected in only 26.85% of cells (**Supplementary Table 2**). Compared to the cell type composition of the whole dataset, the sets of cells removed by the *total probe counts* and *control probe ratio* filters show a much higher fraction of non-malignant cells (**Supplementary Figure 13C**). In contrast, the *area* filter removed non-malignant cells at a frequency consistent with their overall prevalence; however, among these discarded cells some cell types are more (e.g., TAMs) or less (e.g., T cells) represented compared to the non-malignant fraction of the whole dataset. Notably, the application of the *combined* filtering strategy mitigated these compositional biases.

To directly compare *combined* and QS-based filters, we applied two thresholds to the SpaceTrooper quality score: a threshold chosen to remove the same number of cells as the *combined* filter (named as *QS equivalent*; QS threshold = 0.340) and the 3MAD-based cutoff defined in the previous paragraph (named as *QS ≥ 3MAD*; QS threshold = 0.635). For an equivalent number of removed cells (i.e., 3,185), the *combined* and the *QS equivalent* filters exhibited a similar behavior on all cell types (**Supplementary Figure 13D**). However, their overlap was only partial (IoU = 0.349). *QS ≥ 3MAD*, which removes a higher number of cells (8,404 vs 3,185), captures 89.51% of the cells removed by the *combined* filter, while additionally identifying 5,558 cells (**Supplementary Figure 13E**). Despite differences in overlap and numerosity, the sets of cells discarded by the three methods displayed similar cell-type compositions, with a reduced fraction of breast cancer (BC) cells compared to the full dataset (**Supplementary Figure 13F**).

Given these observations, we asked if the *combined* filter had a different impact on the downstream analysis (kNN graph, UMAP embedding, and cluster computation) compared with what we previously observed in **Figure 2**. Compared to the UMAP embedding obtained without applying any filter (**Figures 2D**), the distribution of cell types in the UMAP embedding obtained after applying the *combined* filter appears visually cleaner (**Supplementary Figure 13G**). Nevertheless, one cluster (cluster 14), dispersed across the embedding, still comprises a mixture of all cell lineages (**Supplementary Figure 13G-H**). Consistently, mean homophily scores confirmed that the *combined* filter retains cells that negatively impact downstream analyses (**Supplementary Figure 13I**). Together, these results indicate that conventional QC metrics, even when applied in combination, are insufficient to fully clean the dataset and ensure reliable downstream analyses. In contrast, SpaceTrooper QS-based approach, with a threshold set on the distribution of the quality score (e.g., *QS ≥ 3MAD*), effectively identifies low-quality cells, thereby removing confounding elements.

### SpaceTrooper quality score is generalizable across gene panel coverage, spatial transcriptomics platforms, and tissue types

SpaceTrooper QS is robust and generalizable across gene-panel depths and imaging-based spatial omics platforms. Application of the workflow to the whole-transcriptome CosMx human pancreas dataset demonstrates that the quality score remains stable even when transcriptomic coverage is dramatically expanded (**Supplementary Figures 7A**, **14A** and **15A**). In this whole-transcriptome setting, QS distributions were consistent across cell types (**Supplementary Figures 14B** and **15B**), indicating that the metric is independent of the number of profiled genes and of lineage-specific expression richness. Notably, most cells excluded by NanoString prior to cell-type annotation overlapped with low-QS cells. Spatial maps of the four QS components closely recapitulated the patterns observed in the CosMx 1k breast cancer dataset (**Supplementary Figures 14C-F** and **15C-F**). Each component highlighted analogous spatial artifacts and tissue features, such as proximity to FOV borders or tissue discontinuities, and their integration into the composite QS yielded coherent and biologically interpretable quality landscapes. Together, these results demonstrate that the QS formulation applies to datasets obtained using gene panels of different size and captures technical artifacts even in datasets with substantially expanded transcriptomic coverage.

To evaluate cross-platform generalizability, we next applied SpaceTrooper to a Xenium human lung cancer dataset. Because Xenium images do not exhibit border discontinuities (see **Supplementary Note 1**), QS computation relied only on cell-size and signal-based components; background signal was automatically excluded during model fitting because fewer than 0.1% of all cells were identified as low-quality examples by this metric (13 cells, 0.008% of total; **Supplementary Table 2**). As in the CosMx breast cancer sample, QS values displayed spatial heterogeneity (**Figure 3A**) without any systematic bias toward specific phenotypes, except for two sparsely represented cell populations (7 pulmonary neuroendocrine and 10 mesothelial cells; **Supplementary Figure 16A**). Although low-quality cells were broadly distributed, two regions showed prominent local enrichment, one within a peripheral vessel and the other in an isolated tissue fragment (boxed regions i and ii in **Figure 3A**). Within the vessel, cells consistently exhibited reduced QS values and low signal density (boxed region i; **Figure 3B**), except for a discrete high-quality area at the vessel edge (boxed region iii; **Figure 3C**). Inspection of the composite fluorescence image (**Supplementary Figure 16B**) revealed technical artifacts underlying these patterns, including out-of-focus areas and substantial variation in cytoplasmic and membrane staining intensities across vessel subregions (**Figures 3B-C**). These morphological inconsistencies impaired segmentation, leading to increased transcript misassignment outside cell boundaries (**Supplementary Figure 16C**; left) and to a higher frequency of cells segmented using nuclear-expansion, a fallback method triggered only after the failure of segmentations based on boundary stain and interior RNA^46^ (**Supplementary Figure 16C**; right). All such cells were correctly flagged by SpaceTrooper as low quality (**Figures 3B-C**). A second problematic area, the detached tissue patch at the top of the slide (boxed region ii; **Supplementary Figure 16B**), showed low signal density and QS values, together with impaired staining intensity relatively to the rest of the tissue, again suggesting segmentation inaccuracies (**Supplementary Figure 16D**). Finally, SpaceTrooper marked as low-quality some spatial multiplets (boxed region iv in **Supplementary Figure 16B**, and **Supplementary Figure 16E**).

**Figure 3.**
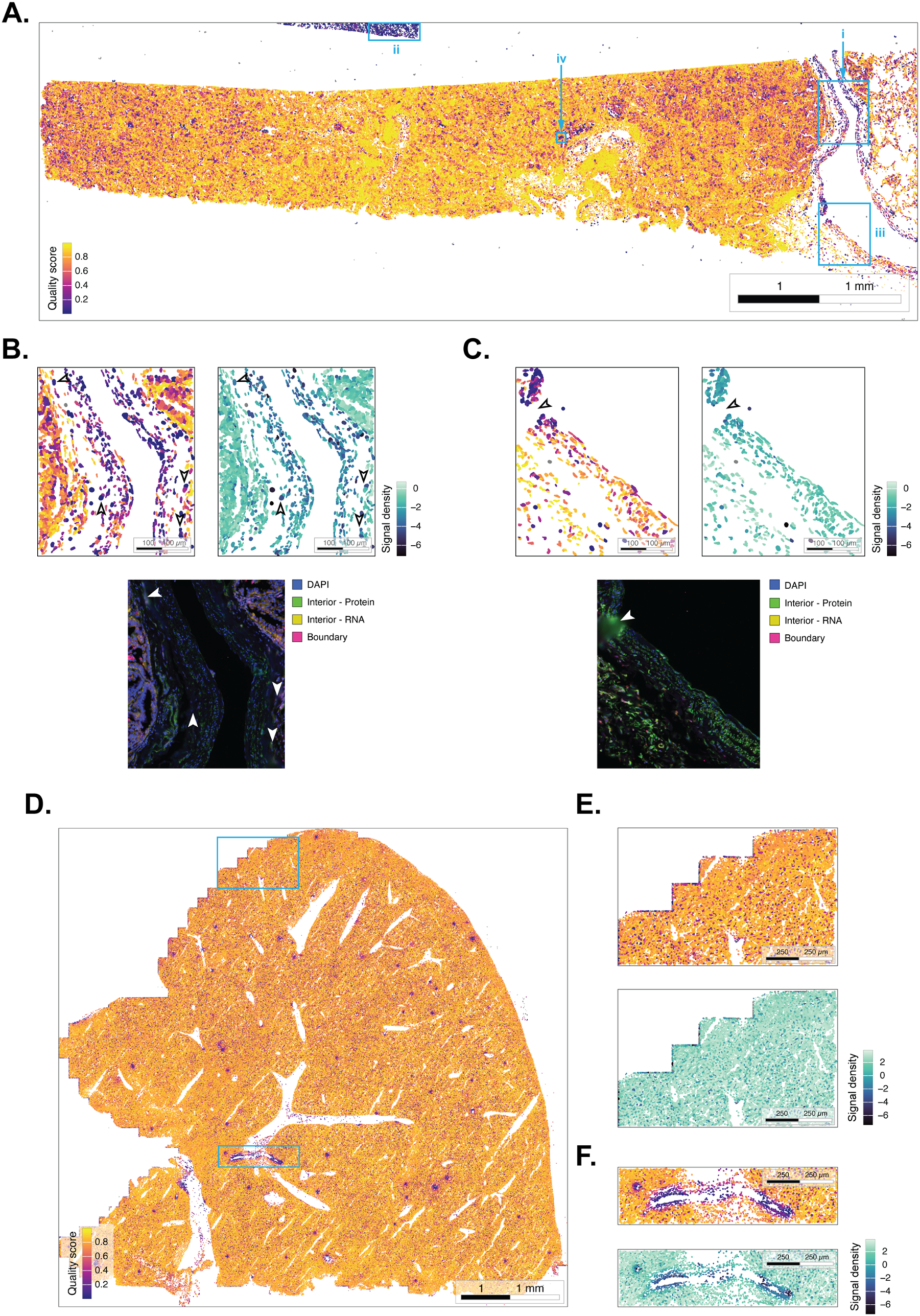
SpaceTrooper QS is generalizable across spatial transcriptomics platforms. **A.** Spatial map of the quality score across the Xenium human lung cancer dataset, with highlighted two regions within a peripheral vessel (boxed regions i and iii), one region in an isolated tissue fragment (boxed region ii), and one region within the tissue mass (boxed region iv). **B.** Spatial distribution of the QS, signal density, and composite fluorescence signals in boxed region i. White arrows indicate technical artifacts as out-of-focus regions, uneven cytoplasmic (interior protein and RNA) and membrane (boundary) staining, and segmentation inaccuracies. **C.** Same as in **B** for the area highlighted in boxed region iii. **D.** Spatial distribution of the quality score in the Vizgen MERFISH mouse liver dataset. Light blue boxes indicate examples of peripheral and interior regions comprising low-QS cells. **E.** Magnified view of a peripheral low-QS region located at the left side of the image. Reduced QS in this area (top) is primarily driven by diminished signal density (bottom). **F.** Same as in **E** for an interior low-QS region. In panels **D**, **E**, and **F**, cells shown in grey correspond to cells with zero counts.

SpaceTrooper generalizability across imaging-based spatial omics technologies and tissue types was further evaluated using the Vizgen MERFISH mouse liver dataset. Consistent with Xenium data, MERFISH does not exhibit systematic FOV border-related shape distortions; therefore, the QS formulation excluded the aspect ratio component. In addition, the cell size component was automatically discarded during model fitting due to the near absence of outliers (3 cells, corresponding to 0.00076% of the dataset). The spatial distribution of QS values across the entire tissue section revealed some regions with low quality cells (**Figure 3D**). Quality score values were consistent across cell types (**Supplementary Figure 17A**). One low-QS region is located at the left side of the image and coincides with an incompletely acquired tissue boundary. Component-level inspection indicates that reduced QS in this area is primarily driven by diminished signal density (**Figure 3E**), consistent with an external border effect affecting transcript detection (**Supplementary Figure 17B**). Regions of low-QS are observed also within the interior tissue and are again predominantly driven by reduced signal density (**Figure 3F**). In contrast to the peripheral artifact, here the pattern is associated with tissue areas exhibiting a depletion of hepatocytes in their cellular composition (**Supplementary Figure 17C**).

Collectively, these results demonstrate that SpaceTrooper identifies low quality cells, accounting for diverse artifacts, across marker panels of varying complexity, multiple spatial transcriptomics platforms, and diverse tissue types and species.

### The quality score is generalizable across modalities

We next evaluated the applicability of SpaceTrooper to a different molecular modality by applying the workflow to a publicly available human tonsil dataset generated using a panel of protein markers. As observed in previous analyses, the quality score (**Figure 4A**) was comparable across cell types (**Supplementary Figure 18A**). Spatial mapping of QS values revealed several localized clusters of low-quality cells, including the region highlighted in **Figure 4B**. In protein-based assays, background signal is a particularly critical factor, as antibody imaging is susceptible to non-specific binding that can confound marker interpretation. In the highlighted region, a compact central group of cells exhibited markedly elevated background signal (**Figure 4C**), while the remaining QS components contributed minimally to the overall score (**Supplementary Figure 18B-D**), indicating that background noise was the primary driver of the reduced QS. Closer examination of marker expression in this area revealed that, within a central vessel, identified by the SMA fluorescence signal (white in **Figure 4D**) and expression (**Figure 4E**), there are objects lacking a detectable nucleus (**Figure 4D** and **Figure 4F**) that simultaneously express mutually incompatible markers. These include keratins and CD68, (**Figures 4D**, 4G, and 4H), as well as markers of T (ICOS in **Figure 4D** and FOXP3 in **Supplementary Figure 18E**) and epithelial (**Supplementary Figure 18F**) cells. The morphology and spatial location of these cells are consistent with erythrocytes, but this interpretation cannot be directly validated due to the absence of erythrocyte-specific markers in the protein panel. Since erythrocytes are known to exhibit autofluorescence^6,47^, this property likely underlies the observed anomalous expression profiles. Accordingly, the elevated background signal and marker misassignment support their classification as low-quality cells. All in all, these results demonstrate that the QS effectively captures modality-specific sources of technical noise and reliably identifies low-quality objects across both RNA- and protein-based spatial omics.

**Figure 4.**
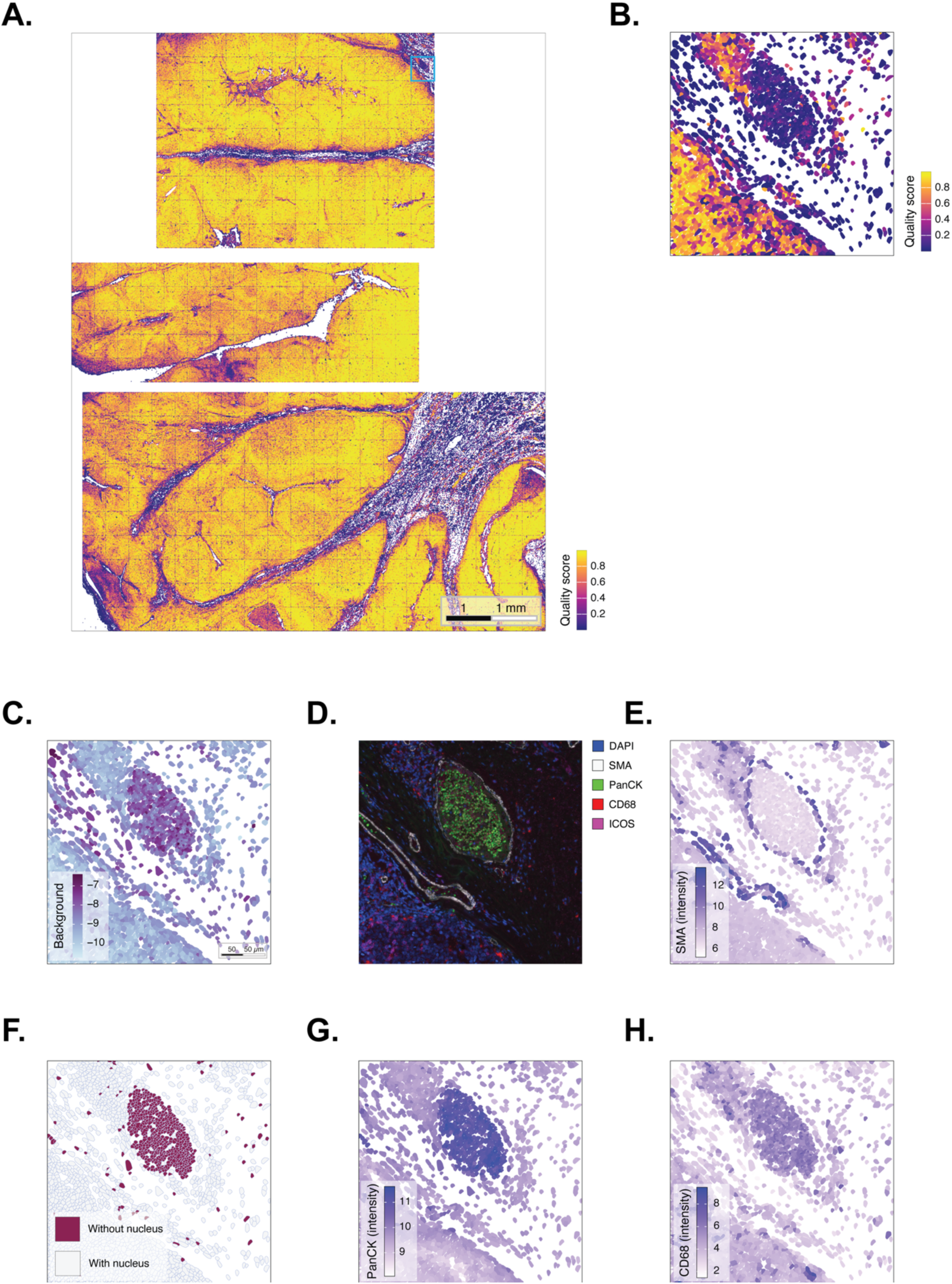
SpaceTrooper QS is generalizable across molecular modalities and captures protein-specific artifacts. **A.** Spatial map of the quality score across the CosMx human tonsil protein dataset. **B.** Magnified region highlighting a localized group of low-QS cells (in light blue in panel **A**). **C.** Background component of the QS in the region of panel **B**, revealing a compact central group of cells with elevated background signal. **D.** Composite fluorescence image of the same region, with DAPI in blue, SMA in white, PanCK in green, CD68 in red, and ICOS in magenta. **E.** Spatial distribution of SMA log_2_-transformed intensity. **F.** Spatial distribution of cells with null nucleus area in the region of panel **B**. **G.** Spatial distribution of PanCK log_2_-transformed intensity. **H.** Spatial distribution of CD68 log_2_-transformed intensity.

## DISCUSSION

Imaging-based single-cell spatial omics technologies have transformed our ability to interrogate tissue organization by linking molecular profiles to precise spatial context. As these technologies mature, data preprocessing, and quality control (QC) in particular, has emerged as a critical determinant of the reliability, reproducibility, and interpretability of downstream analyses. The rapid diversification of platforms, molecular modalities, spatial resolutions, and tissue contexts has exposed fundamental limitations of fixed threshold-based QC strategies largely inherited from dissociation-based single-cell sequencing workflows. Therefore, there is a growing need for QC frameworks that are quantitative, reproducible, and scalable across datasets while avoiding reliance on predefined thresholds.

Existing QC approaches for spatial omics span a broad methodological spectrum, yet no consensus framework has emerged for systematic, cell level quality assessment in imaging-based single cell spatial omics^25–31^. Low-quality cells arising from tissue defects, segmentation errors^42,44,45,48–50^, or experimental artifacts are typically identified through visual inspection, fixed threshold, or modality-specific metrics^26–28,31^. While commonly used metrics such as cell size, total counts, or control probe proportions can be informative, they are strongly influenced by tissue biology, panel design, and experimental conditions, and often fail to capture complex technical artifacts. As a result, QC decisions based on individual metrics risk overlooking subtle but systematic sources of technical bias.

SpaceTrooper addresses these limitations by introducing a unified, data-driven quality score (QS) for imaging-based spatial omics data. Rather than evaluating individual QC metrics in isolation, SpaceTrooper integrates complementary components that capture cell morphology, signal density, background signal, and spatial distortions into a single, interpretable score. This integration enables the detection of combinations of artifacts, such as segmentation errors coupled with signal loss or elevated background, that are difficult to identify using single-metric approaches. Importantly, the QS is comparable across cell types of the various datasets, ensuring that cells are flagged based on technical characteristics rather than lineage-specific molecular abundance. This distinction is particularly critical in spatial omics, where RNA and protein levels can vary widely due to intrinsic biological heterogeneity, tissue integrity, and section quality.

A defining feature of SpaceTrooper is its adaptive, data-driven formulation. Rather than relying on predefined thresholds or platform-specific heuristics, the QS is learned directly from the empirical distributions of QC components within each dataset using adaptive outlier detection and regularized logistic regression. This design allows the score to adjust to differences in tissue type, molecular modality, panel size, and platform-specific artifacts, while remaining interpretable and comparable across datasets. Consistent with this formulation, SpaceTrooper identified diverse technical artifacts across datasets, including segmentation errors, histopathological features, out-of-focus imaging regions, tissue detachment, and FOV border artifacts. Notably, the QS captured not only isolated low-quality cells but also spatially coherent regions of reduced quality, reflecting localized experimental or tissue-related failures. The component-level structure of the QS further provides insight into the specific factors driving quality degradation, an essential advantage in imaging-based spatial omics where understanding the origin of artifacts can inform both filtering strategies and experimental optimization.

Critically, quality differences identified by SpaceTrooper negatively impacted downstream analyses. In the CosMx breast cancer dataset, low-QS cells formed an artifact-driven mixed cluster characterized by reduced cell-type confidence and homogeneity. Removing these cells using a data-adaptive, QS-based threshold substantially improved cluster coherence without introducing cell-type biases. In contrast, conventional QC strategies based on combination of fixed-threshold filters failed to fully resolve artifactually mixed cell populations.

A central challenge for spatial omics QC is generalizability across technologies, resolutions, and molecular modalities. SpaceTrooper addresses this challenge by focusing on QC components shared across imaging-based platforms, while allowing platform-specific features, such as FOV border distortions, to be incorporated only when relevant. Analyses across targeted and whole-transcriptome CosMx datasets, Xenium RNA data, MERFISH data, and CosMx protein assays demonstrate that the QS remains stable across gene panel sizes, RNA and protein modalities, and diverse tissue types from different species, supporting its broad applicability.

In summary, SpaceTrooper provides a robust, adaptable, and interpretable QC framework that addresses key shortcomings of current spatial omics preprocessing strategies. By integrating multiple sources of technical variation into a single data-driven score and embedding QC within established analytical infrastructures, SpaceTrooper provides a practical foundation for standardized, scalable, and reproducible quality control in imaging-based single-cell spatial omics.

## METHODS

### SpaceTrooper computational framework

#### Definition of the cell quality score

We defined a per-cell quality score (QS) that quantifies cell quality by integrating multiple metrics that capture diverse sources of technical artifacts. To this end, we systematically examined the cell metadata provided by NanoString CosMx, 10x Genomics Xenium, and Vizgen MERFISH in a set of publicly available datasets, processed with the standard pipeline provided by the manufacturers. These metadata include morphological characteristics of cells and nuclei, measures of molecular content and control probe signal, spatial attributes relative to the field of view, and fluorescence signals (**Supplementary Table 1**). From the metrics shared across the three technologies, we selected *cell size* (either cell *area* or cell *volume*) to represent cell morphology and capture potential segmentation artifacts; the *signal density*, defined as the log2-transformed ratio of total probe counts to the cell size for RNA platforms, and as the log2-transformed sum of the average intensity for each protein in the cell for protein assays, to quantify the overall molecular content; the *proportion of control probe counts*, defined as the log2-transformed ratio of total control probe counts to total probe counts, to estimate the background noise; and the *aspect ratio*, defined as the ratio of cell width to height, to identify artifacts such as fractured or distorted cells close to the field of view (FOV) border in the standard NanoString CosMx pipeline (**Supplementary Note 1**). To quantify the background signal, we used negative control probes as well as blank controls, depending on their availability across technologies^30^. For NanoString CosMx datasets, the aspect ratio was computed as the ratio between a cell projection along the x-axis and its projection along the y-axis.

For each cell *k*, the quality score *QS_k_* is defined as the logarithm of the odds (logit) that cell *k* is of high- versus low-quality, given a set of *n* predictor components:

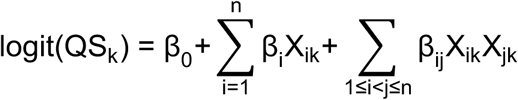

where *n = 3* for 10x Genomics Xenium and Vizgen MERFISH (which generate composite datasets through spatial overlap of FOVs) and *n = 4* for NanoString CosMx (which processes each FOV independently). The predictor components are defined as follows: *X_1k_* is cell area for CosMx and Xenium and cell volume for MERFISH; *X_2k_* is the signal density calculated as described above; *X_3k_* is the background signal quantified as described above; and *X_4k_=* |*log_2_*(*aspect ratio_k_*)|*·*𝕀(*dist.border_k_<dist.thr*) is defined only for NanoString CosMx data to account for artifacts in cells located near the FOV borders (*border effect*). Here, *dist.border_k_* denotes the distance between the centroid of cell *k* and the nearest FOV border and *dist.thr* is the threshold distance used to determine whether a cell is considered close to the border. This threshold can be defined as either a fixed value (e.g., 50 pixels) or adaptively estimated from the data, for instance, as the radius derived from the average cell area under the assumption of circular cell shapes. The term X_ik_X_jk_ denotes the interaction between each pair of predictor components, and the β coefficients are estimated from the data during model fitting.

### Quantification of the cell quality score

The β_0_, β_i_ and β_ij_ coefficients are estimated by fitting a penalized generalized linear model (GLM) using the *glmnet* R package (v4.1-10)^51^. The model is fitted using a binomial family with a logistic link function, which models the probability of each cell being of high quality as a sigmoid transformation of the linear predictor. To mitigate overfitting and improve generalization, we apply ridge regularization, which shrinks towards zero the magnitude of coefficient estimates while retaining all predictor components in the model. The optimal regularization parameter (λ) is selected via k-fold cross-validation as implemented in the *cv.glmnet* function, ensuring a data-driven balance between model bias and variance. We chose ridge regularization over lasso penalty because the predictor components might be partially correlated and biologically interdependent (e.g., cell area and probe count often covary; see **Supplementary Figure 1** and **Supplementary Note 1**), making coefficient shrinkage preferable to variable selection. In the fitting phase, for each dataset, a set of cells is constructed to include an equal number of low- and high-quality cells. For each component, low- and high-quality cells are defined as outliers and central values, respectively, based on the component’s empirical distribution (**Supplementary Table 2**). Outlier detection is performed using either the Medcouple (MC) method^35,36^, implemented in the *mc* function of the *robustbase* R package (v0.99-6), or the Median Absolute Deviation (MAD) method, implemented in the *isOutlier* function from the *scuttle* Bioconductor package (v1.18.0)^52^. The appropriate method is selected based on the symmetry of the component distribution, quantified using the *skewness* function from the *e1071* R package (v1.7-16). Predictor components with significant asymmetry (nonzero skewness) are processed with the MC-based approach, whereas approximately symmetric predictors are analyzed using the MAD-based method. For cell size, low-quality cells are defined as those with a size exceeding either the Medcouple-adjusted upper boundary or the median plus 3 MAD (**Supplementary Figure 1** and **4**). For signal density, low-quality cells are those with a signal density below the Medcouple-adjusted lower boundary or median minus 3 MAD or, if these thresholds are smaller than the observed minimum, below the first percentile of the signal density distribution (**Supplementary Figure 2** and **4**). For the background signal component, low-quality cells are those with a background signal exceeding either the Medcouple-adjusted upper boundary or median plus 3 MAD (**Supplementary Figure 3** and **4**). Finally, for the border effect (in Nanostring CosMx only), low-quality cells are those with the aspect ratio component below and above the Medcouple-adjusted or the median ± 3 MAD thresholds and with a centroid located within a fixed distance (e.g., 50 pixels; see **Supplementary Figures 5** **and 6**) from a FOV border. Cells classified as low-quality for at least one component are included as low-quality examples in the fitting phase. High-quality examples are defined as cells that are not flagged as low-quality by any component and meet additional selection criteria. Specifically, high-quality cells are required to either have a size (area or volume) within the interquartile range of its distribution or a signal density between the 90th and 99th percentiles of the corresponding distribution. For NanoString CosMx data, high-quality cells are also those having an aspect ratio within the interquartile range of its distribution and a centroid located at least a fixed distance away (e.g., 50 pixels) from the nearest FOV border. The size, background signal, and border effect components are excluded from the quality score model if fewer than 0.1% of all cells are classified as low-quality for that component. Although our approach uses the data twice, first to define positive and negative examples and then to estimate the GLM coefficients, we observed no indication of overfitting in practice.

### Computational implementation

SpaceTrooper is implemented as an R package within the Bioconductor framework. The workflow for the quantification of the QS begins by generating a standardized *SpatialExperiment* (SPE) object that harmonizes data from CosMx, Xenium, and MERFISH imaging-based spatial omics technologies. SPE objects are constructed by importing the gene expression matrix, cell metadata, cell boundary polygons, and, when available, FOV positions^34^. These data can be read from comma-separated values (*.csv*), HDF5 (*.h5*), Parquet (*.parquet*), or compressed (*.csv.gz*) files using SpaceTrooper’s high-level reader functions. These readers are built upon the import utilities provided by the *SpatialExperimentIO* Bioconductor package (v1.0.0), which includes dedicated parsers for the different technologies. During SPE creation, any metric required for the quantification of the quality-score, but not present in the input, is automatically computed from the imported cells. The labels of cell metadata are also standardized to enforce consistent naming conventions across platforms and assay modalities. Cell quality metrics, as signal density, the proportion of control probe counts, and (for NanoString CosMx data) the distance between each cell centroid and the nearest FOV border, are quantified from SPE cell metadata using SpaceTrooper’s *spatialPerCellQC* function. For spatial data handling and visualization, SpaceTrooper exploits functions of the *sf* R package (v1.0-23), a widely used tool for managing geometric objects such as cell boundary polygons.

## Datasets

### Nanostring CosMX datasets

#### Human breast cancer dataset

The CosMx 1k breast cancer dataset comprises 45 FOVs obtained from a single tissue section of human ductal carcinoma in situ (DCIS). The dataset includes 59,284 segmented cells profiled for 1,000 RNA target genes and 10 negative control probes from the CosMx Universal Cell Characterization Panel. Raw data, including count matrix, cell annotation metadata, transcript counts and spatial coordinates, cell boundary polygons, global FOV positions, and segmentation images, have been downloaded from https://kero.hgc.jp/Breast_Cancer_Spatial.html. Composite fluorescent images were generated FOV by FOV starting from the original raw morphological channels. Each channel was independently contrast-enhanced in Fiji using the *Enhance Contrast* tool. The five resulting grayscale images were then merged according to the following color assignment: 1) DAPI (nuclear staining) in blue, 2) PanCK (epithelial staining) in green, 3) B2M (membrane staining) in yellow, 4) CD68 (macrophage staining) in magenta, and 5) CD45 (immune cell staining) in red.

#### Human pancreas dataset

The CosMx human pancreas dataset comprises 18 FOVs generated from a formalin-fixed paraffin-embedded (FFPE) human pancreas tissue using a pre-commercial version of the Nanostring CosMx Human Whole Transcriptome panel. The dataset includes 48,944 segmented cells profiled for 18,946 RNA target genes, 50 negative control probes, and 2,735 system control codes. The raw data, including count matrix, cell annotation metadata, transcript counts and spatial coordinates, cell boundary polygons, global FOV positions, segmentation images, and InSituType cell type annotations, were downloaded from https://nanostring.com/products/cosmx-spatial-molecular-imager/ffpe-dataset/cosmx-smi-human-pancreas-ffpe-dataset/. Cell type assignments are available for 47,270 cells as NanoString excluded 1,674 cells from InSituType phenotyping (likely using a fixed threshold <300 on total probe counts).

#### Human tonsil dataset

The CosMx human tonsil dataset comprises 383 FOVs generated from a human tonsil FFPE tissue with the 64-plex CosMx Human Immuno-Oncology Protein Panel. The dataset comprises 1,464,610 segmented cells profiled for 62 protein targets, 2 non-specific antibodies (mouse IgG1 and rabbit IgG), and 5 morphological channels. The raw data, including count matrix, cell annotation metadata, cell boundary polygons, global FOV positions, segmentation images, and cell type annotations, were downloaded from https://nanostring.com/products/cosmx-spatial-molecular-imager/ffpe-dataset/cosmx-human-tonsil-ffpe-protein-dataset/. Cell type assignments were provided by NanoString, based on an unsupervised clustering analysis performed with Seurat, followed by manual cluster annotation. Cell type labels are available for 1,244,918 cells. NanoString excluded 219,692 cells from cell phenotyping and labeled these excluded cells as *Dropped*. In addition, we excluded 144 cells from our analysis since 117 cells had polygons with fewer than four vertices, which are incompatible with the *sf_polygon* function of the *sfheaders* (v0.4.5) package that requires at least four vertices to close the polygon ring, and 28 cells lacked information on cell boundaries.

### 10X Genomics Xenium dataset

#### Human lung cancer dataset

The Xenium human lung cancer dataset includes 162,254 cells from an adult human lung adenocarcinoma FFPE tissue. The section has been profiled using the Xenium Human Multi-Tissue and Cancer Panel comprising 377 RNA target genes, 20 negative control probes, 41 negative control code words, and 103 blank code words. The raw data, including count matrix, cell annotation metadata, and cell boundary polygons, were downloaded from https://www.10xgenomics.com/datasets/preview-data-ffpe-human-lung-cancer-with-xenium-multimodal-cell-segmentation-1-standard.

### Vizgen MERFISH dataset

*Mouse liver dataset*. The MERFISH mouse liver dataset includes 395,215 cells from a mouse liver tissue (animal 1, replicate 1) profiled using a gene panel containing 347 RNA target genes and 38 blank controls. The raw data, including gene counts per cell matrix, cell metadata, and cell boundary polygons, were downloaded from https://info.vizgen.com/mouse-liver-data?submissionGuid=01f8b3c0-46a8-414c-92fc-4334f920dfe4. Since MERFISH data includes z-stacked cell polygons, we used the middle plane in spatial visialization to maintain consistency with the representations of cell boundary polygon used for CosMx and Xenium data.

## Data analysis

### Detection of potential segmentation artifacts

To identify potential segmentation artifacts, we applied FastReseg and StarDist to the CosMx 1k breast cancer dataset. FastReseg is an algorithm designed to refine image-based cell segmentation and to detect spatial doublets arising from cell proximity or partial overlap in 2D images^37^. We used FastReseg (v1.0.0; https://github.com/Nanostring-Biostats/FastReseg) with default parameters from the input generation to the *fastReseg_flag_all_errors* function, as described in the package vignette. Cell type annotations, generated by InSituType, were provided as reference labels. For each transcript, FastReseg calculates a score that quantifies the probability that the transcript is consistent with the cell type of its assigned cell. The algorithm then evaluates the spatial dependency of these transcript-level scores within each cell using the *lrtest_nlog10P* metric; elevated values indicate potential segmentation artifacts. Spatial multiplets were identified based on the *lrtest_nlog10P* metric using the default threshold of 5. StarDist is a deep learning-based segmentation method that represents object shapes as star-convex polygons, enabling accurate delineation of overlapping and irregularly shaped cells or nuclei^53^. We applied the StarDist plugin for Fiji (https://imagej.net/plugins/stardist) to DAPI morphological images from FOVs 11 and 12 of the CosMx 1k breast cancer dataset. Before segmentation, raw TIFF images were preprocessed by automatic contrast enhancement (pixel saturation set to 0.35%, with normalization). StarDist was then run using the *2D Versatile* (*fluorescent nuclei*) pretrained model with default parameters (v0.3.0).

### Cell phenotyping

Cell type annotation was performed using InSituType^54^ and scRNA-seq data obtained from cell populations of human and mouse tissues as references^40^ to perform the label transfer onto CosMx, Xenium, and MERFISH datasets. InSituType models the transcript counts for each gene in each cell using a negative binomial distribution and applies an Expectation-Maximization (EM) algorithm to assign cells to cell types while iteratively updating the expected expression profiles for each type^54^. For the CosMx 1k breast cancer dataset, the reference matrix of expected cell type expression profiles was generated using cells derived from an integrated scRNA-seq dataset composed of two DCIS samples^55^ (HTA12_250_4000 and HTA12_247_4000) obtained from the Human Tumor Atlas Network (https://humantumoratlas.org)^56^. Individual samples were analyzed separately and subsequently integrated using the Seurat workflow (v4.3.0). No cell- or gene-level filtering was applied during preprocessing. Prior to integration, data normalization and cell cycle phase scoring were performed independently for each sample. Following integration, cell cycle phase scores were included as covariates during data scaling. Clustering and nonlinear dimensionality reduction were performed using the first 14 principal components. Clusters identified at a resolution of 0.8 were manually annotated based on established breast cancer cell type markers. Three clusters characterized by low-quality or ambiguous marker expression were excluded from the final reference. The integrated matrix therefore comprised 12 distinct cell types. To generate the custom reference profile matrix, we used the *create_profile_matrix* function from the *SpatialDecon* Bioconductor package (v1.12.0), which computes average gene expression profiles for each cell type from a scRNA-seq count matrix and corresponding cell annotations. We provided the scRNA-seq reference together with the raw expression profiles and selected morphological features of the spatial transcriptomics data as input to the *insitutypeML* function from the *InSituType* R package (v1.0.0; (https://github.com/Nanostring-Biostats/InSituType) in supervised mode. All parameters were kept at their default, except for *insufficient_anchors_thresh* which was set to 9 in the *choose_anchors_from_stats* function.

For the Xenium human lung cancer dataset, we downloaded the lung tissue single cell RNA-seq reference from the NanoString cell profile library (https://github.com/Nanostring-Biostats/CellProfileLibrary/blob/master/Human/Adult/Lung_Control_Adams.RData) and used it to run InSituType in supervised mode with default parameters. Cell type labels returned by *insitutypeML* were simplified using the cell groups annotations provided with the scRNA-seq reference. We subsequently merged dendrocyte and pDC clusters into a single *DC* label and combined T and NK groups into a unified *T/NK* label. Finally, 9 cells with extremely low counts (i.e., 1 and 2) were discarded by InSituType and labeled as *undefined*.

For the MERFISH mouse liver dataset, we downloaded expression data and cell annotations for liver mouse cells from the Liver Cell Atlas (https://www.livercellatlas.org/download.php; Mouse StSt)^57^. Stromal cells were further stratified using the fibroblast annotation provided in the same repository. To generate the custom reference profile matrix, we used the *create_profile_matrix* function of *SpatialDecon* Bioconductor package (v1.12.0). InSituType was executed in supervised mode with default settings. Finally, we harmonized cell labels renaming Fibroblast 1, Fibroblast 2, and Capsule fibroblasts as *Fibroblast*, T cells and NK cells as *T/NK*, all dendritic cell populations as *DC*, and Monocytes&Monocyte-derived cells as *Myeloid*. Five cells with extremely low counts (i.e., 1) were discarded by InSituType and labeled as *undefined*.

### Downstream analysis of CosMx 1k breast cancer dataset

Prior to downstream analyses, cells with zero raw counts were removed from the CosMx 1k breast cancer dataset. All subsequent analyses, including i) normalization and scaling of the expression matrix; ii) linear dimensionality reduction by Principal Component Analysis (PCA); iii) construction of a k Nearest Neighbor (kNN) graph followed by Louvain clustering; and iv) computation of a two-dimensional UMAP embedding for visualization, were performed using the *Seurat* package (v5.3.1)^58^, following the recommended workflows. To assess the impact on downstream analyses of cell filtering based on QS, fixed thresholds, or combined criteria, cells were excluded before applying the Seurat workflow. Raw count matrices, derived either from the full dataset or from the filtered subsets, were normalized and variance-stabilized using the *SCTransform* function of the Seurat package. Dimensionality reduction was performed by Principal Component Analysis (PCA) using the *RunPCA* function with default parameters. Based on the proportion of variance explained, we retained the first 25 principal components. We constructed a kNN graph in the PCA space using the *FindNeighbors* function with *k* = 6, followed by Louvain community detection using the *FindClusters* function with a resolution of 1. For visualization, UMAP embedding was generated with the *RunUMAP* function and default parameters. Cell phenotypes were assigned using InSituType, as described above. To simplify graphical representations, we collapsed the fine-grained cell-type annotations into broad lineages using the following criteria: we labeled i) mural cells, myoepithelial cells, and cancer-associated fibroblasts (CAFs) as *Stromal*; ii) blood endothelial cells as *Endothelial*; iii) breast cancer (BC) tumor cells as *BC cells*; iv) mixed BC cells/TAMs as *Mixed BC cells TAMs*; and v) all remaining cell types as *Immune*. In the violin and stacked bar plots, clusters were ordered according to their most represented cell type, using the following cell-type sequence: *Mixed*, *T cells*, *NK cells*, *Plasma cells*, *Mast cells*, *DCs*, *TAMs*, *Blood ECs*, *Mural cells*, *CAFs*, *Myoepithelial cells*, *Mixed BC cells TAMs*, and *BC cells*. Clusters were labeled as *Mixed* when no single cell type accounted for at least 30% of the cells.

### Homophily quantification

Homophily quantifies the degree of similarity among connected nodes within a network, describing the tendency of elements with similar attributes to associate. For any cell *v* of the CosMx 1k breast cancer dataset, we quantified the homophily *h(v)* as the fraction of its neighboring cells that share the same phenotype label:

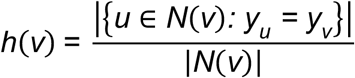

where *N(v)* denotes the set of neighbors of cell *v* in the kNN graph and *y_u_* ∈ {1,…,m} is the phenotype label associated with cell *u*, with *m* being the total number of phenotype labels. We constructed the kNN graph using the *igraph* package (v2.1.4), with each cell represented as a node. Since homophily was computed at the single-cell level, the mean homophily across all cells within a cluster was used to summarize and compare the degree of within-cluster homogeneity.

### Comparison with single-metric thresholding

We selected three widely used quality control (QC) metrics to identify low-quality cells in the CosMx 1k breast cancer dataset using global QC thresholds. Cells were classified as low-quality if they met at least one of the following criteria: i) total detected probes fewer than 20 (total probe count criterion); ii) proportion of control probes greater than 0.01 of the total detected probes (control probe ratio criterion); and iii) cell area greater than 216.14 µm^2^ (area criterion). Thresholds for the total probe count and control probe ratio were adopted from published studies and the NanoString recommended analysis workflow (https://nanostring.com/blog/tips-when-performing-cosmx-data-analysis-with-atomx-sip/#tip-1). The threshold for cell area was obtained using the median plus 3 MAD method^59^. Each QC criterion was evaluated independently and then combined into a unified filtering rule (*combined*), as commonly done in single-cell analysis pipelines. To enable a direct comparison with SpaceTrooper QS filtering using the same number of retained cells, we defined a QS-equivalent threshold (QS = 0.340) that removed the same number of cells as the *combined* approach (3,185 cells; 5.37% of the total). In addition, we defined a conservative QS threshold (QS = 0.635) using the median QS value minus 3 MAD, corresponding to the removal of 8,404 cells (14.18% of the total).

### Computing resources and scalability

All SpaceTrooper analyses were performed on a computer equipped with an Intel Core i9-13900K (3.00 GHz), 32 virtual CPU threads, and 64 GB of RAM. To assess the scalability of the SpaceTrooper workflow, we quantified runtime and memory usage across datasets of increasing cell size (10,000, 20,000, 50,000, 100,000, 200,000, 500,000, and 1,464,610 cells) that were generated randomly subsampling (n = 4 per size) the protein dataset (**Supplementary Figure 19**). Execution time was measured using the *mark* function from the *bench* R package (v1.1.4), while peak memory consumption was estimated using the *peakRAM* function from the *peakRAM* R package (v1.0.2). Measurements were performed both for the complete workflow (**Supplementary Figures 19A-B**) and for the QS quantification step alone (**Supplementary Figures 19C-D**).

## DATA AVAILABILITY

The CosMx 1k breast cancer dataset has been downloaded from https://kero.hgc.jp/Breast_Cancer_Spatial.html. All other data were downloaded directly from Nanostring, 10x Genomics, and Vizgen’s publicly available datasets, including CosMx human pancreas (https://nanostring.com/products/cosmx-spatial-molecular-imager/ffpe-dataset/cosmx-smi-human-pancreas-ffpe-dataset/), CosMx human tonsil (https://nanostring.com/products/cosmx-spatial-molecular-imager/ffpe-dataset/cosmx-human-tonsil-ffpe-protein-dataset/), Xenium human lung cancer (https://www.10xgenomics.com/datasets/preview-data-ffpe-human-lung-cancer-with-xenium-multimodal-cell-segmentation-1-standard), and MERFISH mouse liver

(https://info.vizgen.com/mouse-liver-data?submissionGuid=01f8b3c0-46a8-414c-92fc-4334f920dfe4). The metadata of the spatial omics datasets analyzed in this manuscript and the scRNA-seq reference matrices used for cell phenotyping of the CosMx 1k breast cancer and MERFISH mouse liver samples are publicly available from GitHub (https://github.com/bicciatolab/bicciatolab_data).

## CODE AVAILABILITY

SpaceTrooper is available at Bioconductor (https://bioconductor.org/packages/SpaceTrooper/).

## ACKNOWLEDGMENTS

We thank Helena L. Crowell and Michelangelo Cordenonsi for insightful discussions on imaging-based spatial omics data quality, technical biases, and the molecular characteristics of the analyzed tissues. This work is supported by Fondazione AIRC under 5 per mille 2019 - ID. 22759 program to S.B. and by the European Research Council (ERC) Grant CoG 101171662 to D.Ris.

## AUTHOR CONTRIBUTIONS

BB developed the quality score methodology. BB and MM implemented key package functions and conducted the analyses across all datasets. DRig and BB developed the SpaceTrooper package, including implementation of key functions and adaptation of the code to support Bioconductor classes. All authors contributed to the development of the SpaceTrooper framework and its application to the datasets. OR and MF overviewed methodological development, software implementation, and data analysis. MF, DRis, and SB supervised the overall project. All authors participated in manuscript writing and reviewed and approved the final version.

## Notes

### Competing Interest Statement

The authors have declared no competing interest.

## REFERENCES

1 Asp, M., Bergenstrahle, J. & Lundeberg, J. Spatially Resolved Transcriptomes-Next Generation Tools for Tissue Exploration. Bioessays 42, e1900221 (2020). 10.1002/bies.201900221

2 Moses, L. & Pachter, L. Museum of spatial transcriptomics. Nat Methods 19, 534–546 (2022). 10.1038/s41592-022-01409-2

3 Park, J. et al. Spatial omics technologies at multimodal and single cell/subcellular level. Genome Biol 23, 256 (2022). 10.1186/s13059-022-02824-6

4 Moffitt, J. R., Lundberg, E. & Heyn, H. The emerging landscape of spatial profiling technologies. Nat Rev Genet 23, 741–759 (2022). 10.1038/s41576-022-00515-3

5 Bressan, D., Battistoni, G. & Hannon, G. J. The dawn of spatial omics. Science 381, eabq4964 (2023). 10.1126/science.abq4964

6 Hickey, J. W. et al. Spatial mapping of protein composition and tissue organization: a primer for multiplexed antibody-based imaging. Nat Methods 19, 284–295 (2022). 10.1038/s41592-021-01316-y

7 Rao, A., Barkley, D., Franca, G. S. & Yanai, I. Exploring tissue architecture using spatial transcriptomics. Nature 596, 211–220 (2021). 10.1038/s41586-021-03634-9

8 Palla, G., Fischer, D. S., Regev, A. & Theis, F. J. Spatial components of molecular tissue biology. Nat Biotechnol 40, 308–318 (2022). 10.1038/s41587-021-01182-1

9 Tian, L., Chen, F. & Macosko, E. Z. The expanding vistas of spatial transcriptomics. Nat Biotechnol 41, 773–782 (2023). 10.1038/s41587-022-01448-2

10 Williams, C. G., Lee, H. J., Asatsuma, T., Vento-Tormo, R. & Haque, A. An introduction to spatial transcriptomics for biomedical research. Genome Med 14, 68 (2022). 10.1186/s13073-022-01075-1

11 Wu, Y., Cheng, Y., Wang, X., Fan, J. & Gao, Q. Spatial omics: Navigating to the golden era of cancer research. Clin Transl Med 12, e696 (2022). 10.1002/ctm2.696

12 Walsh, L. A. & Quail, D. F. Decoding the tumor microenvironment with spatial technologies. Nat Immunol 24, 1982–1993 (2023). 10.1038/s41590-023-01678-9

13 Bollhagen, A. & Bodenmiller, B. Highly Multiplexed Tissue Imaging in Precision Oncology and Translational Cancer Research. Cancer Discovery 14, 2071–2088 (2024). 10.1158/2159-8290.CD-23-1165

14 Lazar, E. & Lundeberg, J. Spatial architecture of development and disease. Nat Rev Genet (2025). 10.1038/s41576-025-00892-5

15 Satija, R., Farrell, J. A., Gennert, D., Schier, A. F. & Regev, A. Spatial reconstruction of single-cell gene expression data. Nat Biotechnol 33, 495–502 (2015). 10.1038/nbt.3192

16 Caicedo, J. C. et al. Data-analysis strategies for image-based cell profiling. Nature Methods 14, 849–863 (2017). 10.1038/nmeth.4397

17 Dries, R. et al. Giotto: a toolbox for integrative analysis and visualization of spatial expression data. Genome Biol 22, 78 (2021). 10.1186/s13059-021-02286-2

18 Dries, R. et al. Advances in spatial transcriptomic data analysis. Genome Res 31, 1706–1718 (2021). 10.1101/gr.275224.121

19 Atta, L. & Fan, J. Computational challenges and opportunities in spatially resolved transcriptomic data analysis. Nat Commun 12, 5283 (2021). 10.1038/s41467-021-25557-9

20 Palla, G. et al. Squidpy: a scalable framework for spatial omics analysis. Nat Methods 19, 171–178 (2022). 10.1038/s41592-021-01358-2

21 Moses, L. et al. Voyager: exploratory single-cell genomics data analysis with geospatial statistics. bioRxiv (2023). 10.1101/2023.07.20.549945

22 Heumos, L. et al. Best practices for single-cell analysis across modalities. Nature Reviews Genetics 24, 550–572 (2023). 10.1038/s41576-023-00586-w

23 Fang, S. et al. Computational Approaches and Challenges in Spatial Transcriptomics. Genomics Proteomics Bioinformatics 21, 24 (2023). 10.1016/j.gpb.2022.10.001

24 Chen, J. G. et al. Giotto Suite: a multiscale and technology-agnostic spatial multiomics analysis ecosystem. Nat Methods 22, 2052–2064 (2025). 10.1038/s41592-025-02817-w

25 Vierdag, W.-M. A. M. & Saka, S. K. A perspective on FAIR quality control in multiplexed imaging data processing. Frontiers in Bioinformatics Volume 4 - 2024 (2024). 10.3389/fbinf.2024.1336257

26 Kummerfeld, E. et al. Artifacts in spatial transcriptomics data: their detection, importance, prevalence, and prevention. Brief Bioinform 26 (2025). 10.1093/bib/bbaf306

27 Totty, M., Hicks, S. C. & Guo, B. SpotSweeper: spatially aware quality control for spatial transcriptomics. Nat Methods 22, 1520–1530 (2025). 10.1038/s41592-025-02713-3

28 Martin, N. et al. MerQuaCo: a computational tool for quality control in image-based spatial transcriptomics. (2025). 10.7554/elife.105149.1

29 Marco Salas, S., et al. Optimizing Xenium In Situ data utility by quality assessment and best-practice analysis workflows. Nat Methods 22, 813–823 (2025). 10.1038/s41592-025-02617-2

30 Ozirmak Lermi, N., et al. Comparison of imaging based single-cell resolution spatial transcriptomics profiling platforms using formalin-fixed paraffin-embedded tumor samples. Nat Commun 16, 8499 (2025). 10.1038/s41467-025-63414-1

31 Baker, G. J. et al. Quality control for single-cell analysis of high-plex tissue profiles using CyLinter. Nature Methods 21, 2248–2259 (2024). 10.1038/s41592-024-02328-0

32 Chen, K. H., Boettiger, A. N., Moffitt, J. R., Wang, S. & Zhuang, X. RNA imaging. Spatially resolved, highly multiplexed RNA profiling in single cells. Science 348, aaa6090 (2015). 10.1126/science.aaa6090

33 Rozenblatt-Rosen, O. et al. The Human Tumor Atlas Network: Charting Tumor Transitions across Space and Time at Single-Cell Resolution. Cell 181, 236–249 (2020). 10.1016/j.cell.2020.03.053

34 Righelli, D. et al. SpatialExperiment: infrastructure for spatially-resolved transcriptomics data in R using Bioconductor. Bioinformatics 38, 3128–3131 (2022). 10.1093/bioinformatics/btac299

35 Brys, G., Hubert, M. & Struyf, A. A robust measure of skewness. J Comput Graph Stat 13, 996–1017 (2004). Doi 10.1198/106186004x12632

36 Hubert, M. & Vandervieren, E. An adjusted boxplot for skewed distributions. Computational Statistics & Data Analysis 52, 5186–5201 (2008). 10.1016/j.csda.2007.11.008

37 Wu, L., Beechem, J. M. & Danaher, P. Using transcripts to refine image based cell segmentation with FastReseg. Sci Rep 15, 30508 (2025). 10.1038/s41598-025-08733-5

38 Gatenby, R. A. & Gillies, R. J. Why do cancers have high aerobic glycolysis? Nat Rev Cancer 4, 891–899 (2004). 10.1038/nrc1478

39 Shekhar, M. P. et al. Comedo-ductal carcinoma in situ: A paradoxical role for programmed cell death. Cancer Biol Ther 7, 1774–1782 (2008). 10.4161/cbt.7.11.6781

40 WHO Classification of Tumours Editorial Board. Breast Tumours. 5th edn, Vol. 2 (International Agency for Research on Cancer (IARC), 2019).

41 Mehrabi, M., Amini, F. & Mehrabi, S. Active Role of the Necrotic Zone in Desensitization of Hypoxic Macrophages and Regulation of CSC-Fate: A hypothesis. Front Oncol 8, 235 (2018). 10.3389/fonc.2018.00235

42 Stringer, C. & Pachitariu, M. Cellpose3: one-click image restoration for improved cellular segmentation. Nat Methods 22, 592–599 (2025). 10.1038/s41592-025-02595-5

43 Luecken, M. D. & Theis, F. J. Current best practices in single-cell RNA-seq analysis: a tutorial. Mol Syst Biol 15, e8746 (2019). 10.15252/msb.20188746

44 Petukhov, V. et al. Cell segmentation in imaging-based spatial transcriptomics. Nat Biotechnol 40, 345–354 (2022). 10.1038/s41587-021-01044-w

45 Bruhns, M. et al. Effects of segmentation errors on downstream-analysis in highly-multiplexed tissue imaging. PLoS Comput Biol 21, e1013350 (2025). 10.1371/journal.pcbi.1013350

46 Xenium In Situ Multimodal Cell Segmentation: Workflow and Data Highlights. https://www.10xgenomics.com/support/in-situ-gene-expression/documentation/steps/assay/cell-segmentation-tech-note. (10X Genomics, 2025).

47 Whittington, N. C. & Wray, S. Suppression of Red Blood Cell Autofluorescence for Immunocytochemistry on Fixed Embryonic Mouse Tissue. Current Protocols in Neuroscience 81 (2017). 10.1002/cpns.35

48 Mitchel, J., Gao, T., Cole, E., Petukhov, V. & Kharchenko, P. V. Impact of Segmentation Errors in Analysis of Spatial Transcriptomics Data. bioRxiv, 2025.2001.2002.631135 (2025). 10.1101/2025.01.02.631135

49 Chen, H. & Murphy, R. F. Evaluation of cell segmentation methods without reference segmentations. Mol Biol Cell 34, ar50 (2023). 10.1091/mbc.E22-08-0364

50 He, S. et al. High-plex imaging of RNA and proteins at subcellular resolution in fixed tissue by spatial molecular imaging. Nat Biotechnol 40, 1794–1806 (2022). 10.1038/s41587-022-01483-z

51 Tay, J. K., Narasimhan, B. & Hastie, T. Elastic Net Regularization Paths for All Generalized Linear Models. J Stat Softw 106 (2023). 10.18637/jss.v106.i01

52 McCarthy, D. J., Campbell, K. R., Lun, A. T. & Wills, Q. F. Scater: pre-processing, quality control, normalization and visualization of single-cell RNA-seq data in R. Bioinformatics 33, 1179–1186 (2017). 10.1093/bioinformatics/btw777

53 Schmidt, U., Weigert, M., Broaddus, C. & Myers, G. Cell Detection with Star-Convex Polygons. Lect Notes Comput Sc 11071, 265–273 (2018). 10.1007/978-3-030-00934-2_30

54 Danaher, P. et al. Insitutype: likelihood-based cell typing for single cell spatial transcriptomics. bioRxiv, 2022.2010.2019.512902 (2022). 10.1101/2022.10.19.512902

55 Iglesia, M. D. et al. Differential chromatin accessibility and transcriptional dynamics define breast cancer subtypes and their lineages. Nat Cancer 5, 1713–1736 (2024). 10.1038/s43018-024-00773-6

56 de Bruijn, I. et al. Sharing data from the Human Tumor Atlas Network through standards, infrastructure and community engagement. Nat Methods 22, 664–671 (2025). 10.1038/s41592-025-02643-0

57 Guilliams, M. et al. Spatial proteogenomics reveals distinct and evolutionarily conserved hepatic macrophage niches. Cell 185, 379–396.e338 (2022). 10.1016/j.cell.2021.12.018

58 Hao, Y. et al. Dictionary learning for integrative, multimodal and scalable single-cell analysis. Nat Biotechnol 42, 293–304 (2024). 10.1038/s41587-023-01767-y

59 Pitino, E. et al. STAMP: Single-cell transcriptomics analysis and multimodal profiling through imaging. Cell 188, 5100–5117 e5126 (2025). 10.1016/j.cell.2025.05.027

