## Supplementary Figures for "*SpaceTrooper*: a quality control framework for imaging-based spatial omics data"

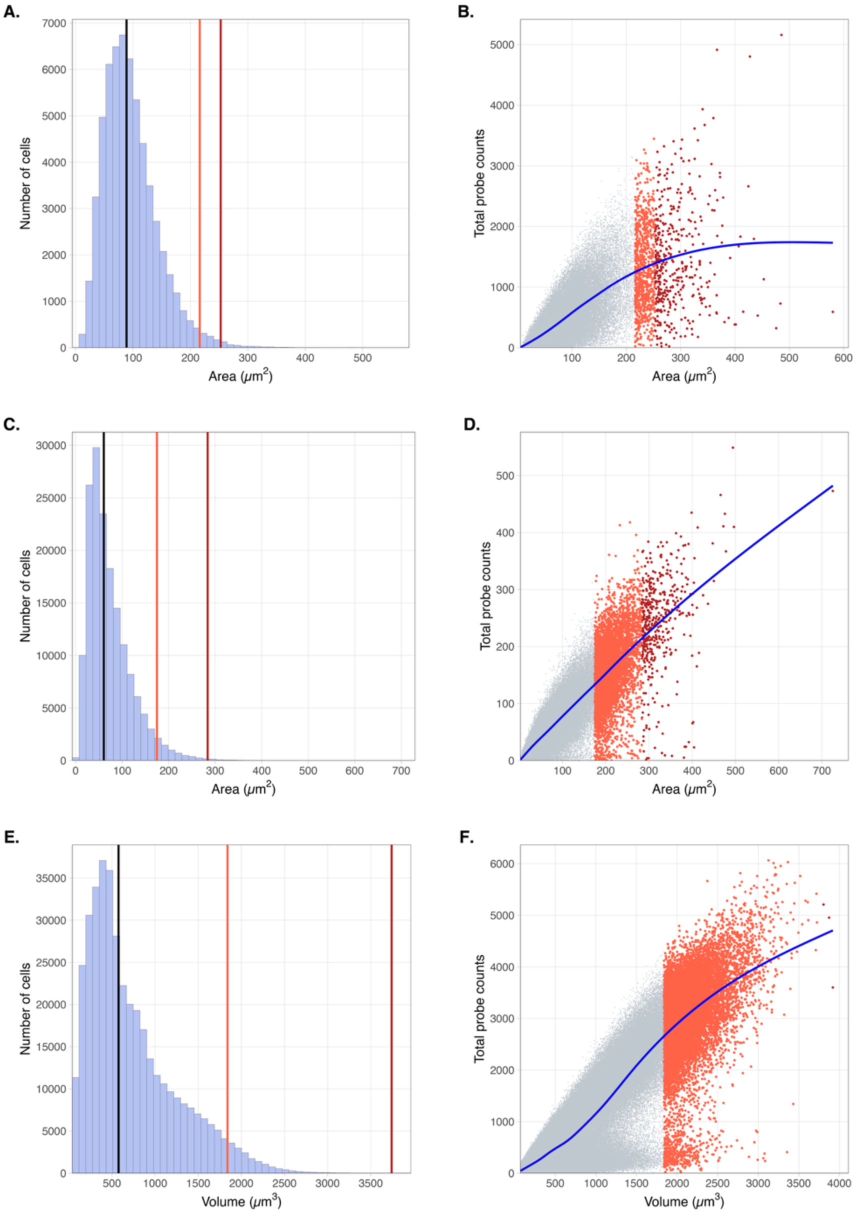


**Supplementary Figure 1. Cell size distributions and their relationship with total transcript counts in the spatial transcriptomics datasets. A.** Distribution of cell area in the CosMx 1k breast cancer dataset. The median, median plus 3 median absolute deviation (3MAD), and Medcouple-adjusted upper boundary are indicated by black, red, and dark red vertical lines, respectively. **B**. Relationship between cell area and the total number of measured mRNA molecules in the CosMx 1k breast cancer dataset. Cells identified as area outliers based on 3MAD and Medcouple thresholds are highlighted in red and dark red, respectively. **C.** Same as in **A** for the Xenium human lung cancer dataset. **D.** Same as in **B** for the Xenium human lung cancer dataset. **E.** Same as in **A** and **C** for the MERFISH mouse liver dataset with cell size represented by cell volume. **F.** Same as in **B** and **D** for the MERFISH mouse liver dataset with cell size represented by cell volume.

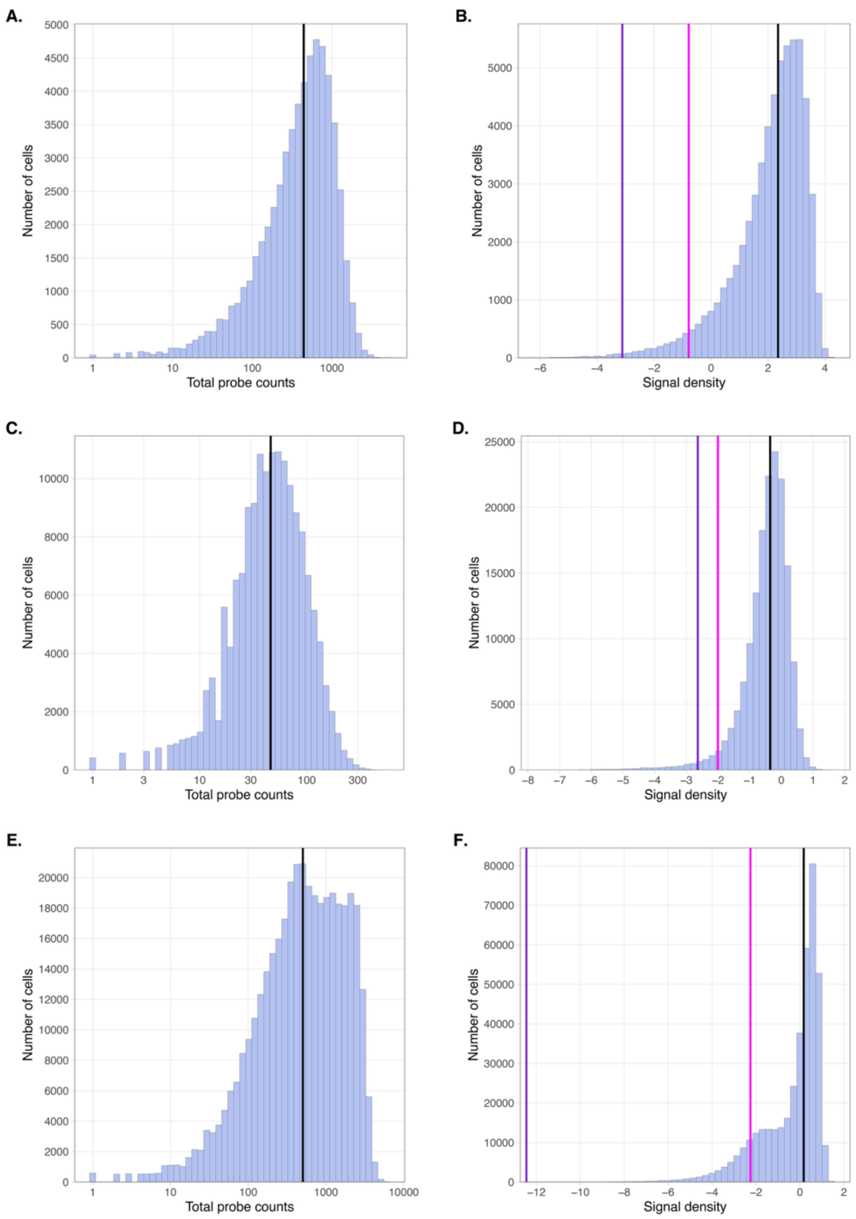


**Supplementary Figure 2. Distribution of total probe counts and signal density in the spatial transcriptomics datasets. A.** Distribution of total probe counts per cell in the CosMx 1k breast cancer dataset. The black vertical line indicates the median total probe count. **B**. Distribution of signal density, defined as the total counts normalized by cell size and $log_2$-transformed, in the CosMx 1k breast cancer dataset. Magenta and violet vertical lines indicate thresholds based on median minus 3 MAD and Medcouple-adjusted statistics, respectively, used to identify low-density outlier cells. **C.** Same as in **A** for the Xenium human lung cancer dataset. **D.** Same as in **B** for the Xenium human lung cancer dataset. **E.** Same as in **A** and **C** for the MERFISH mouse liver dataset. **F.** Same as in **B** and **D** for the MERFISH mouse liver dataset.

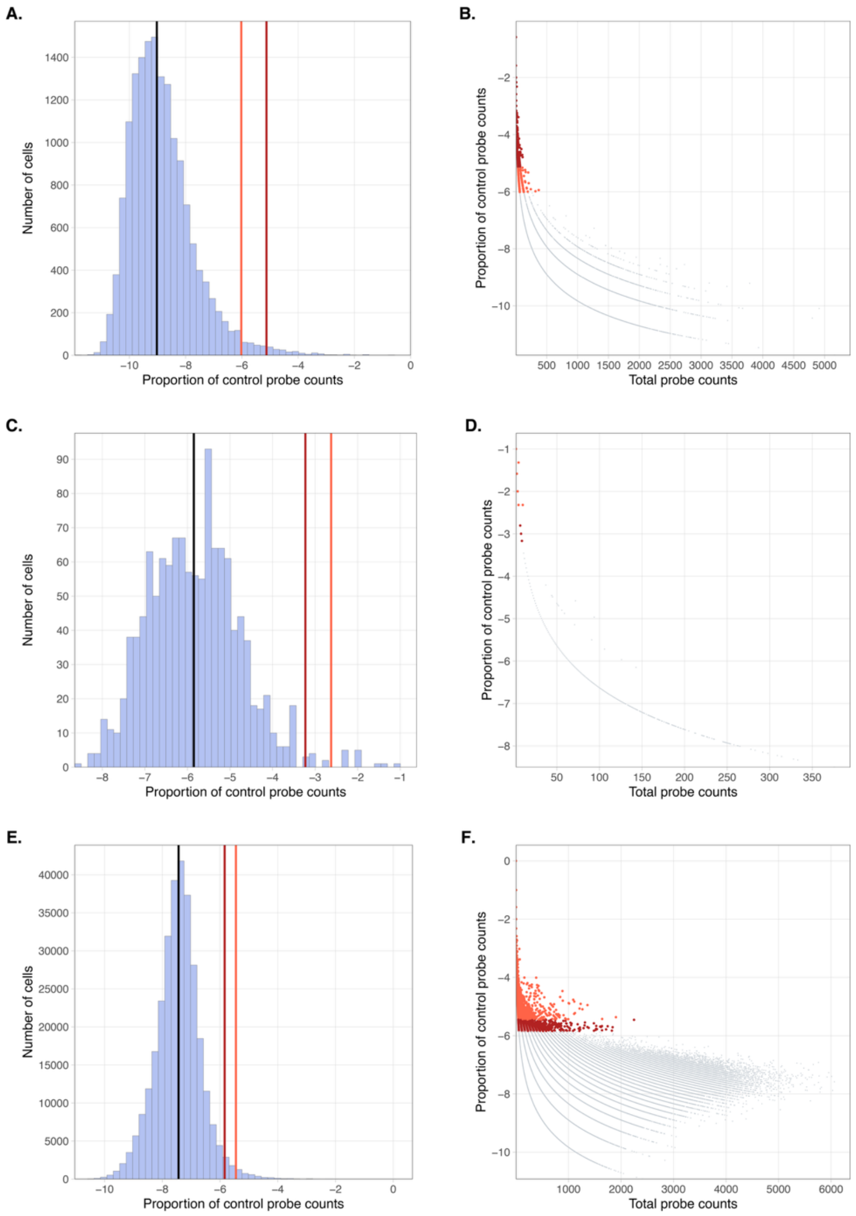


**Supplementary Figure 3. Distributions of control probe signal and its relationship with total transcript counts in the spatial transcriptomics datasets. A.** Distribution of the $log_2$-transformed proportion of control probe counts relative to total counts in the CosMx 1k breast cancer dataset. The black, red, and dark red vertical lines indicate the median, median plus 3 MAD (3MAD), and Medcouple-adjusted upper boundary, respectively. **B**. Relationship between the $log_2$-transformed proportion of control probe counts and total mRNA counts in the CosMx 1k breast cancer dataset. Cells identified as background signal outliers based on 3MAD and Medcouple thresholds are highlighted in red and dark red, respectively. **C.** Same as in **A** for the Xenium human lung cancer dataset. **D.** Same as in **B** for the Xenium human lung cancer dataset. **E.** Same as in **A** and **C** for the MERFISH mouse liver dataset. **F.** Same as in **B** and **D** for the MERFISH mouse liver dataset. Only cells with non-zero control probe counts are considered.

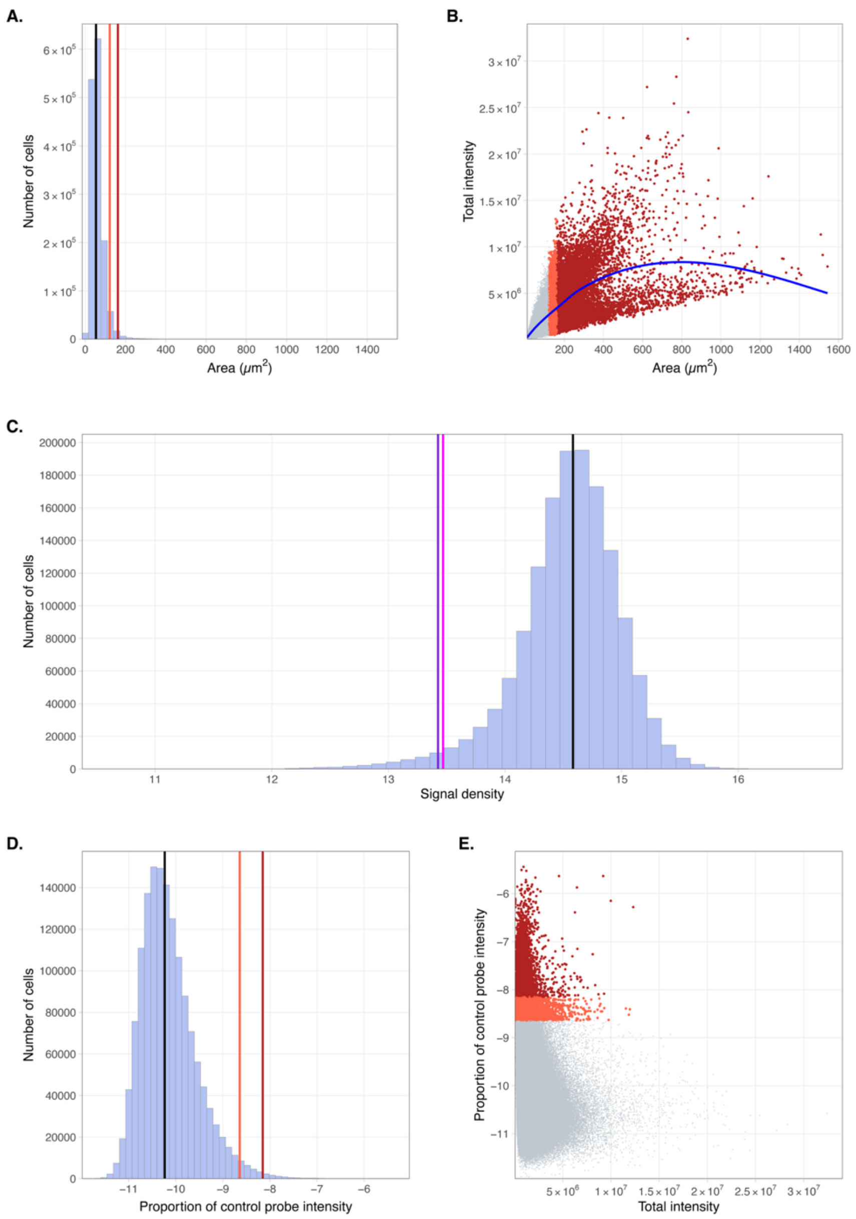


**Supplementary Figure 4. Cell size, signal density, and control probe signal intensity in the CosMx protein human tonsil dataset. A.** Distribution of cell area. The median, median plus 3 MAD (3MAD), and Medcouple-adjusted upper boundary are indicated by black, red, and dark red vertical lines, respectively. **B**. Relationship between cell area and total protein signal intensity, calculated as the sum of the average intensity for each protein in the cell multiplied by cell area. Cells identified as size outliers based on 3MAD and Medcouple thresholds are highlighted in red and dark red, respectively. **C.** Distribution of signal density. Magenta and violet vertical lines indicate thresholds based on 3MAD and Medcouple, respectively. **D.** Distribution of the $\log_2$-transformed proportion of control probe intensity relative to total protein signal intensity. The black, red, and dark red vertical lines indicate the median, 3MAD, and Medcouple-adjusted upper boundary, respectively. **E.** Relationship between the $\log_2$-transformed proportion of control probe intensity and total protein signal intensity. Cells identified as background signal outliers based on 3MAD and Medcouple thresholds are highlighted in red and dark red, respectively.

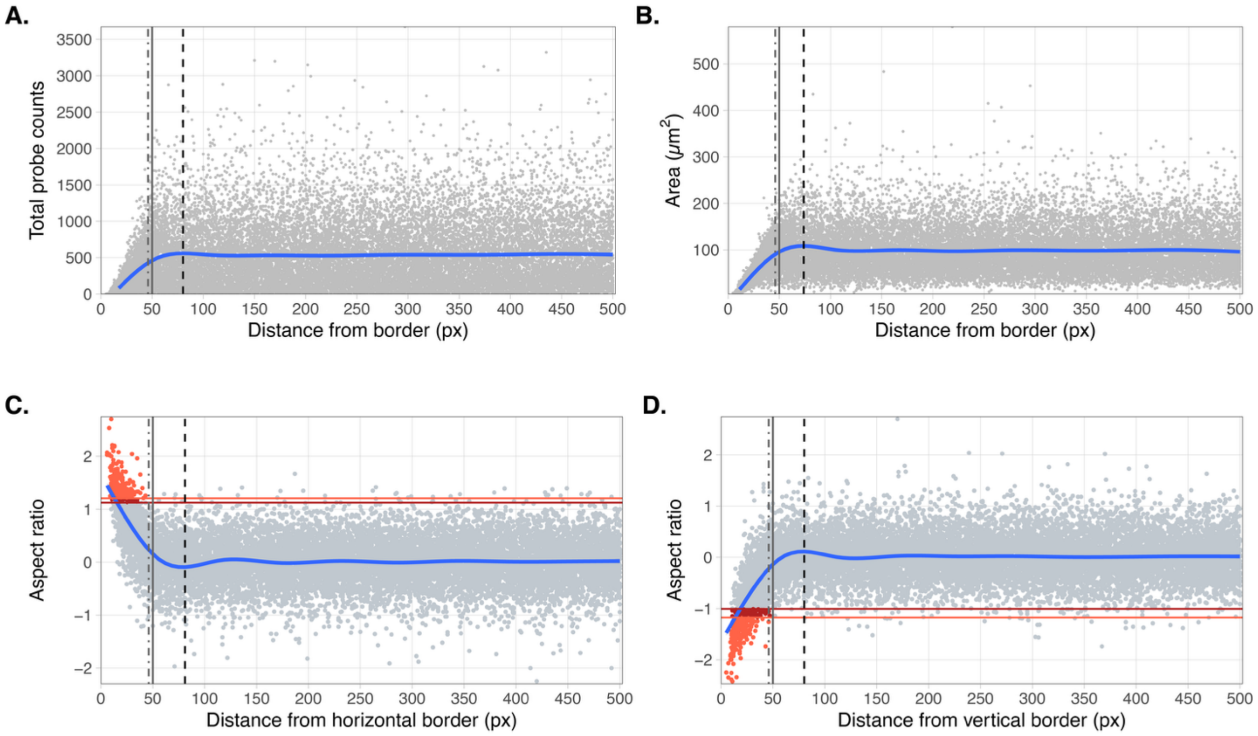


**Supplementary Figure 5. Total probe counts, cell area, and aspect ratio relative to FOV borders in the CosMx 1k breast cancer dataset. A.** Total probe counts plotted against the distance from each cell centroid to the nearest FOV border. **B**. Same as in **A** for the cell area. **C.** $Log_2$-transformed aspect ratio as a function of the distance from each cell centroid to the nearest horizontal FOV border. **D.** $Log_2$-transformed aspect ratio as a function of the distance from each cell centroid to the nearest vertical FOV border. In all panels, the average cell radius, a manually selected threshold of 50 pixels, and the location of the knee of the metric interpolation (blue curve) are indicated by dot-dash, solid, and dashed lines, respectively. Cells identified as aspect ratio outliers based on 3 median absolute deviation and Medcouple-adjusted thresholds are highlighted in red and dark red, respectively.

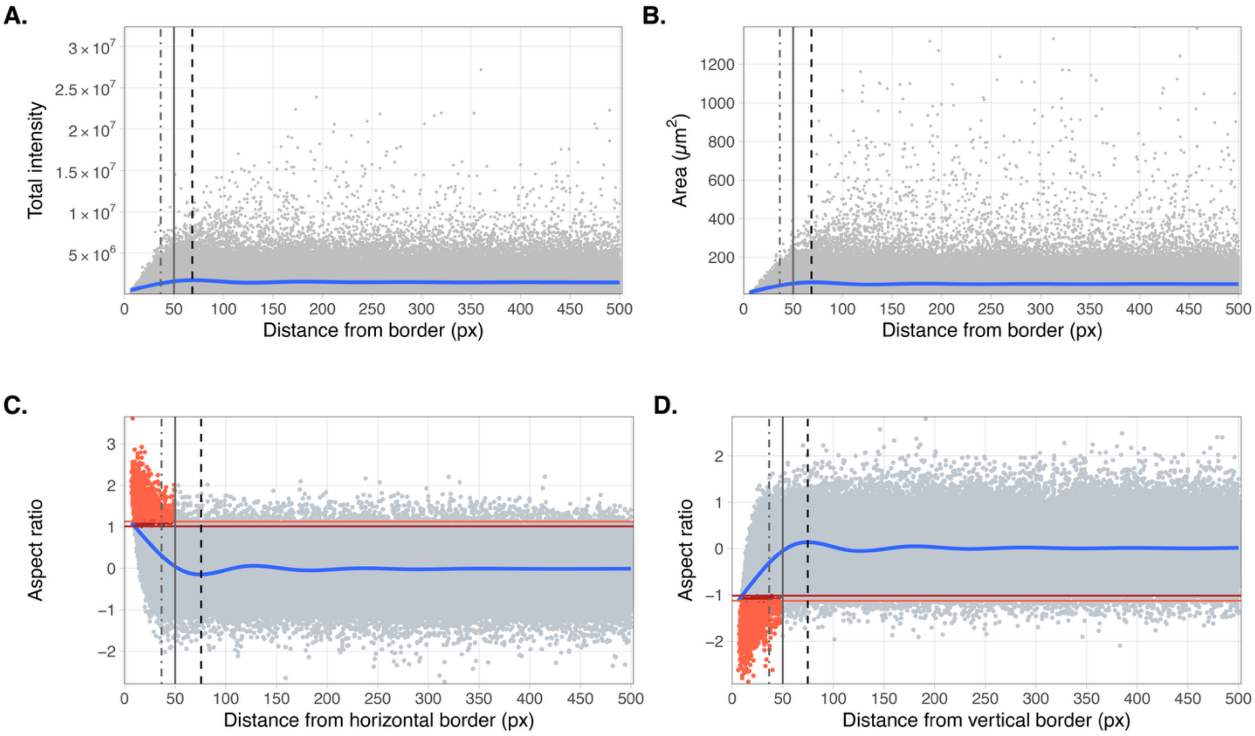


**Supplementary Figure 6. Total probe intensity, area, and cell aspect ratio against distance to FOV borders in the CosMx protein human tonsil dataset. A.** Total intensity plotted against the distance from each cell centroid to the nearest FOV border. **B**. Same as in **A** for the cell area. **C.** Log$_2$ aspect ratio as a function of the distance from each cell centroid to the nearest horizontal FOV border. **D.** Log$_2$ aspect ratio as a function of the distance from each cell centroid to the nearest vertical FOV border. In all panels, the average cell radius, a manually selected threshold of 50 pixels, and the location of the knee of the metric interpolation (blue curve) are indicated by dot-dash, solid, and dashed lines, respectively. Cells identified as aspect ratio outliers based on 3 median absolute deviation and Medcouple-adjusted thresholds are highlighted in red and dark red, respectively.

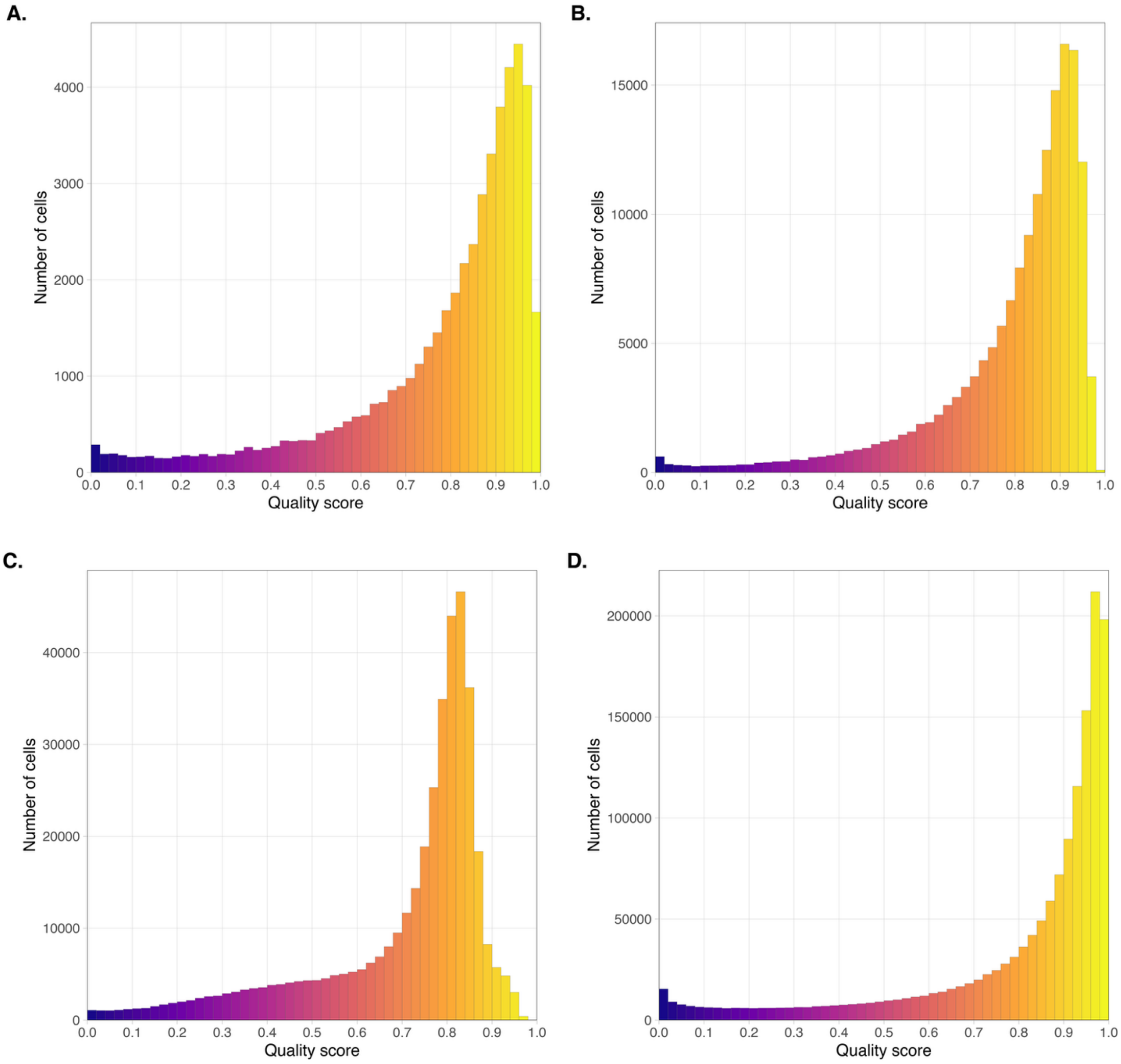


**Supplementary Figure 7. Distribution of the cell quality score in the analyzed datasets. A.** CosMx human pancreas. **B**. Xenium human lung cancer. **C.** MERFISH mouse liver. **D.** CosMx human tonsil (protein assay).

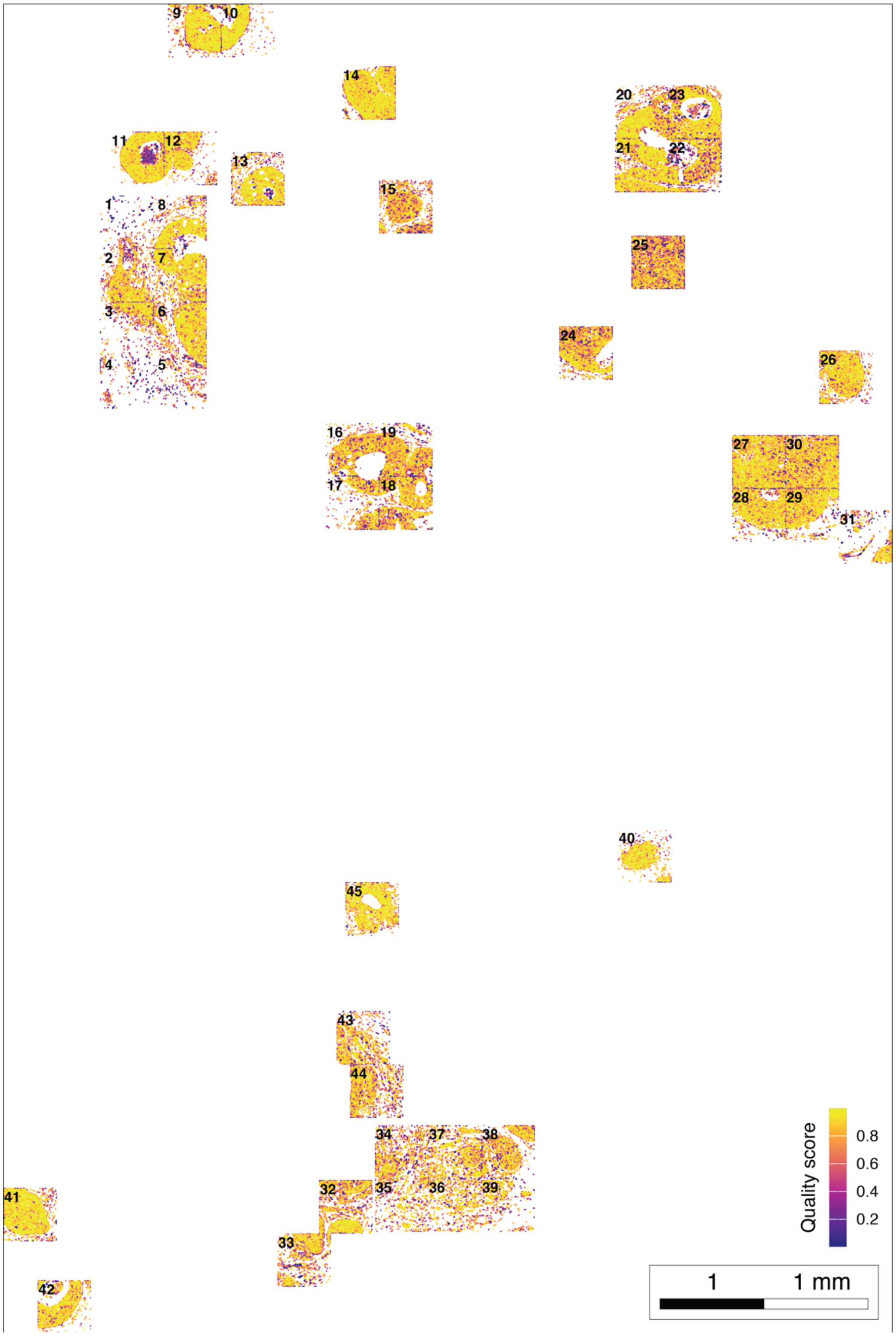


**Supplementary Figure 8. Spatial distribution of cell quality scores in the CosMx 1k breast cancer dataset.** Spatial map of the quality score across the CosMx DCIS breast cancer tissue section. Numbers in the left upper corner of each area indicate the corresponding FOV number.

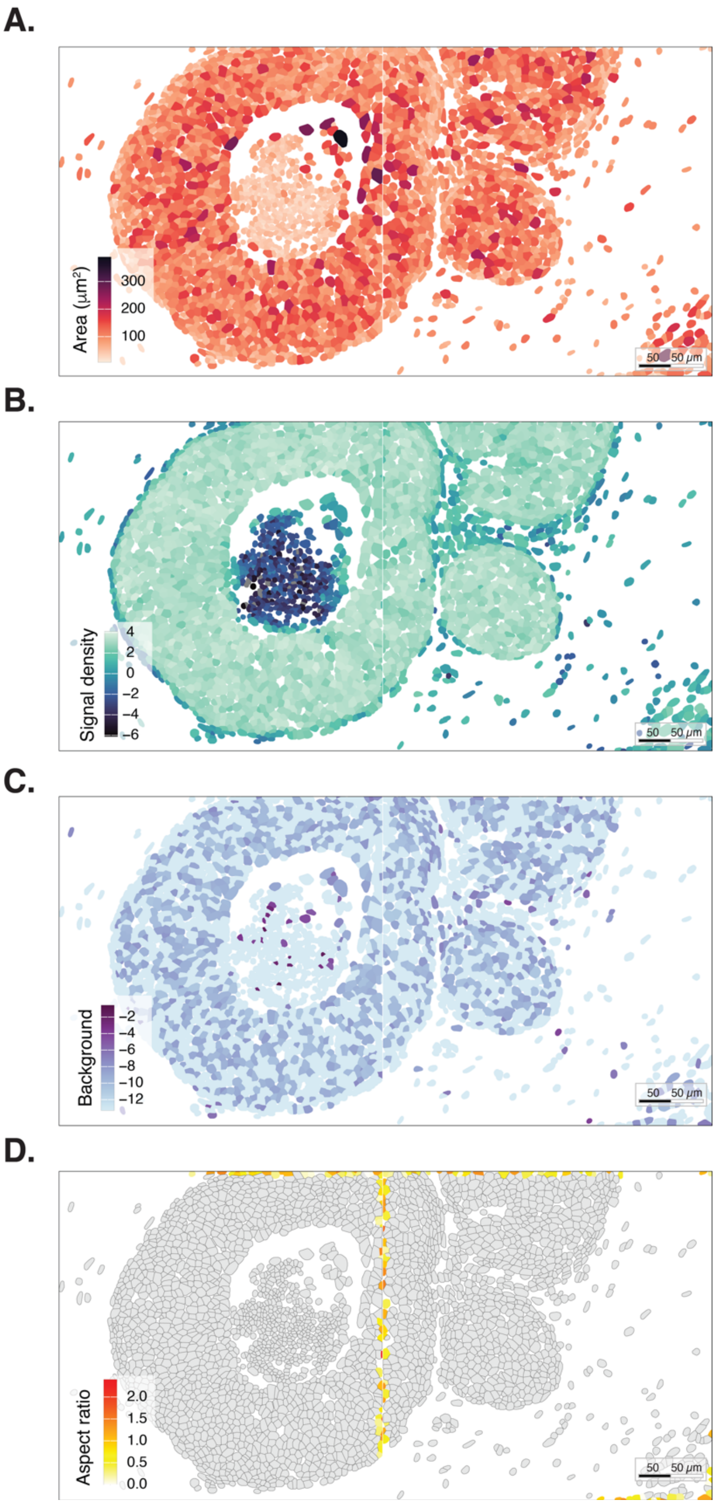


**Supplementary Figure 9. Spatial distribution of the QS components across FOVs 11 and 12 of the CosMx 1k breast cancer dataset. A.** Cell area (in $\mu m^2$). **B.** Signal density. **C.** Background signal. **D.** Absolute $\log_2$ aspect ratio for cells whose centroid lies within 50 px of the nearest FOV boundary.

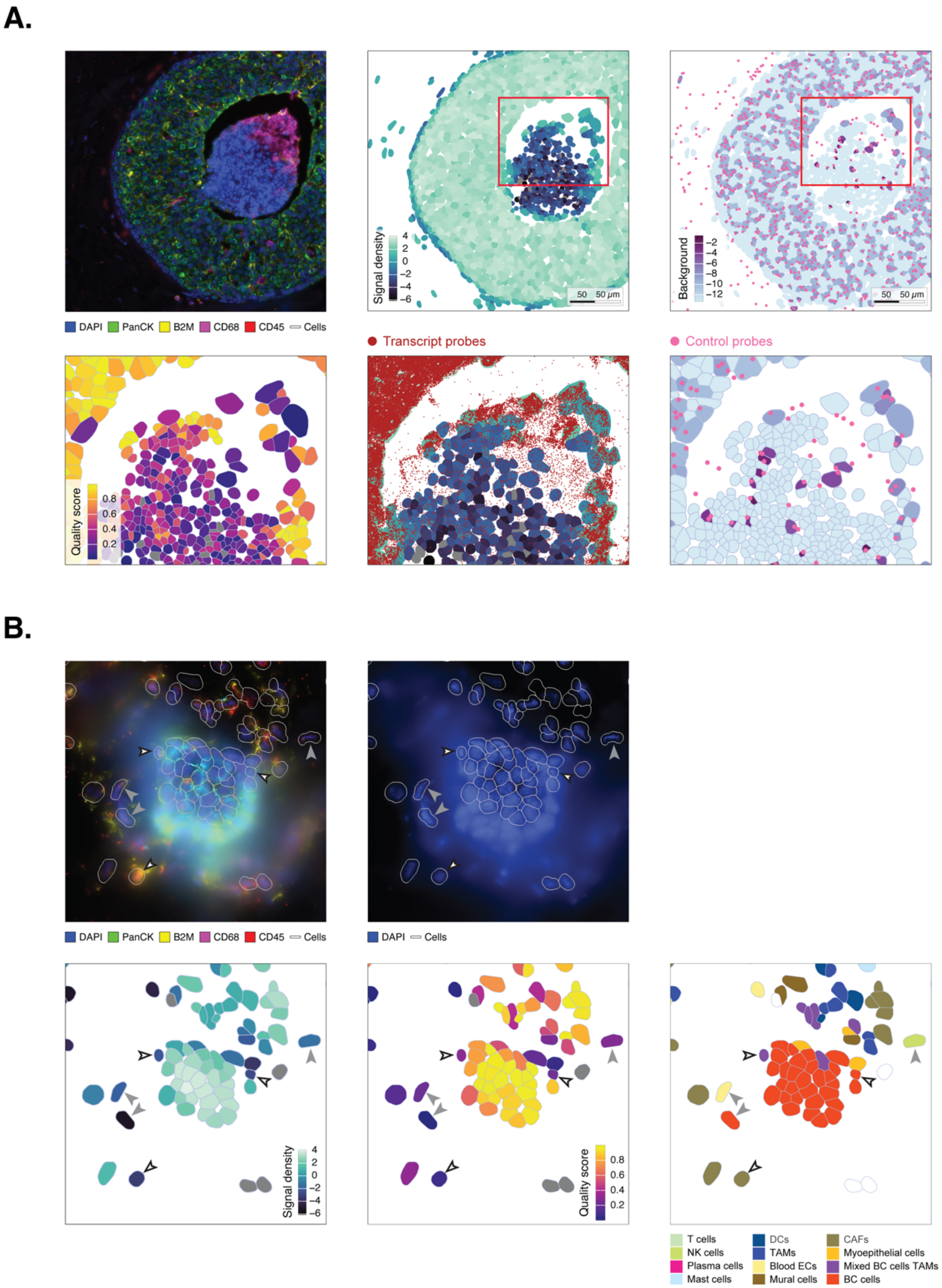


**Supplementary Figure 10. SpaceTrooper identifies artifacts induced by the sample histopathological characteristics and out-of-focus imaging defects. A.** Detection of a comedonecrotic area within the lumen of FOV 11 in the CosMx 1k breast cancer dataset. The composite image (top left) and signal density (middle) reveal a central necrotic core characterized by dense pyknotic DAPI-positive debris, loss of cytokeratin staining, and near-complete depletion of transcript probes (in dark red). Segmented cells within this region show low QS values (bottom left) and elevated background values (right; control probes in magenta), consistent with signal degradation. **B.** Identification of an out-of-focus imaging artifact in FOV 31 of the same dataset. The composite and DAPI images (top) display a blurred region indicative of local tissue detachment and loss of optical focus. Despite being segmented, cells in the out-of-focus area exhibit reduced transcriptional content and are assigned consistently low QS values (bottom right and middle; white-filled arrowheads). Cells surrounding the blurred zone (grey-filled arrows) show morphological abnormalities consistent with mechanical stress and receive low QS values, independently of their phenotypes (bottom right). In the signal density and QS plots, cells shown in grey correspond to cells with zero counts; the same cells are shown in white in the cell type plot.

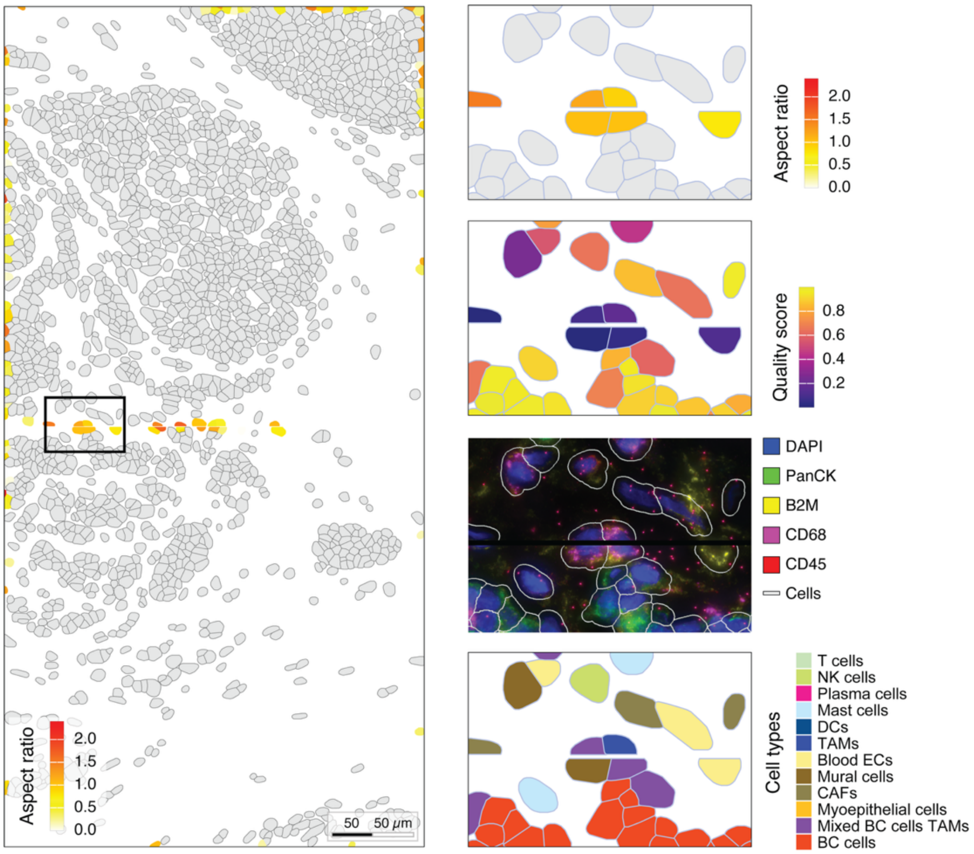


**Supplementary Figure 11. SpaceTrooper identifies artifacts at FOV boundaries.** Spatial mapping of cell aspect ratio (left) and corresponding zoomed region (right) reveal border-associated distortions at the interface between FOV 38 and 39 in the CosMx 1k breast cancer dataset, where duplicated tissue regions lead to double representations of the same cells. SpaceTrooper highlights inconsistencies at FOV borders through abnormal aspect-ratio values and low QS. Although displaying similar shapes and nearly identical patterns in the composite immunofluorescence image (middle right), these duplicated polygons receive inconsistent phenotype assignments across the two FOVs (bottom right).

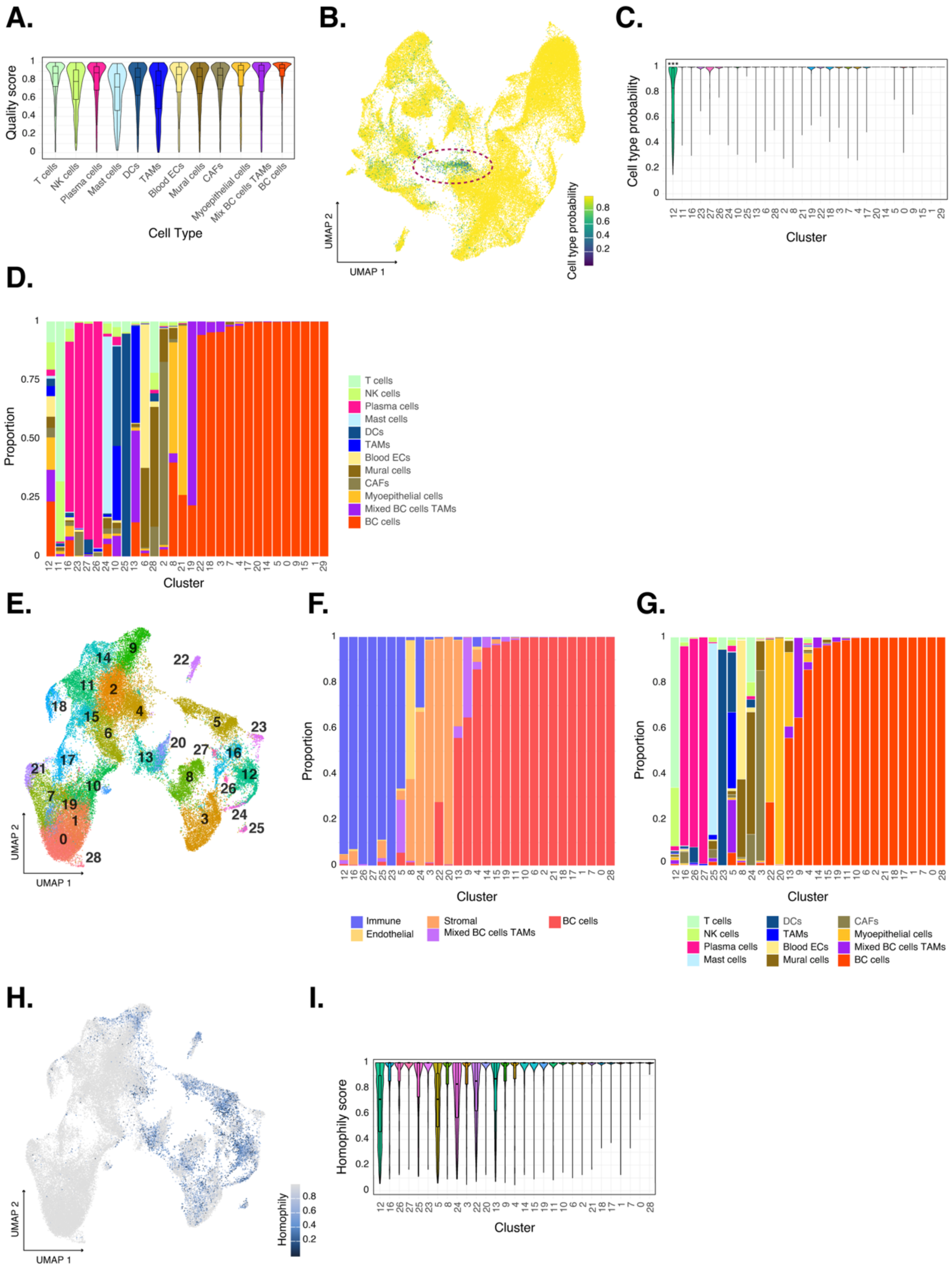

**Supplementary Figure 12. Low-quality cells drive artifactual mixing in UMAP embeddings and affect downstream analyses. A.** Distribution of SpaceTrooper quality scores (QS) across cell types. QS values are broadly comparable among cell types, indicating that the QS is not biased toward specific phenotypes. **B.** UMAP embedding of the full dataset (same as in **Figure 2A**) colored by InSituType cell-type assignment probability. A central region with reduced assignment confidence is highlighted by the dashed ellipse. **C.** Distribution of InSituType assignment probabilities across clusters. Cluster 12 exhibits significantly lower probabilities values compared with each of the other clusters (one-sided Wilcoxon test with Benjamini–Hochberg correction; *** $p<0.001$ for all comparisons). Clusters are ordered as in **Figure 2C**. **D.** Stacked bar plot showing the proportion of annotated cell types per cluster in the full dataset. Cluster 12 displays a marked heterogeneity, with substantial contributions from multiple cell types. Clusters are ordered as in **Figure 2C**. **E.** UMAP embedding (same as in **Figure 2H**) colored by cluster identity, after filtering low-quality cells using the QS 3MAD threshold (median minus 3 MAD). **F.** Stacked bar plot summarizing the proportion of major cell lineages per cluster after filtering low-quality cells using the QS 3MAD threshold. Clusters are ordered according to cell type composition as described in the Method section. **G.** Stacked bar plot showing detailed cell-type composition per cluster after filtering low-quality cells using the QS 3MAD threshold. Clusters are ordered as in **F**. **H.** UMAP embedding (same as in **Figure 2H**) colored by homophily score after filtering low-quality cells using the QS 3MAD threshold. **I.** Distribution of homophily scores across clusters after filtering low-quality cells using the QS 3MAD threshold. Clusters are ordered as in **F**.

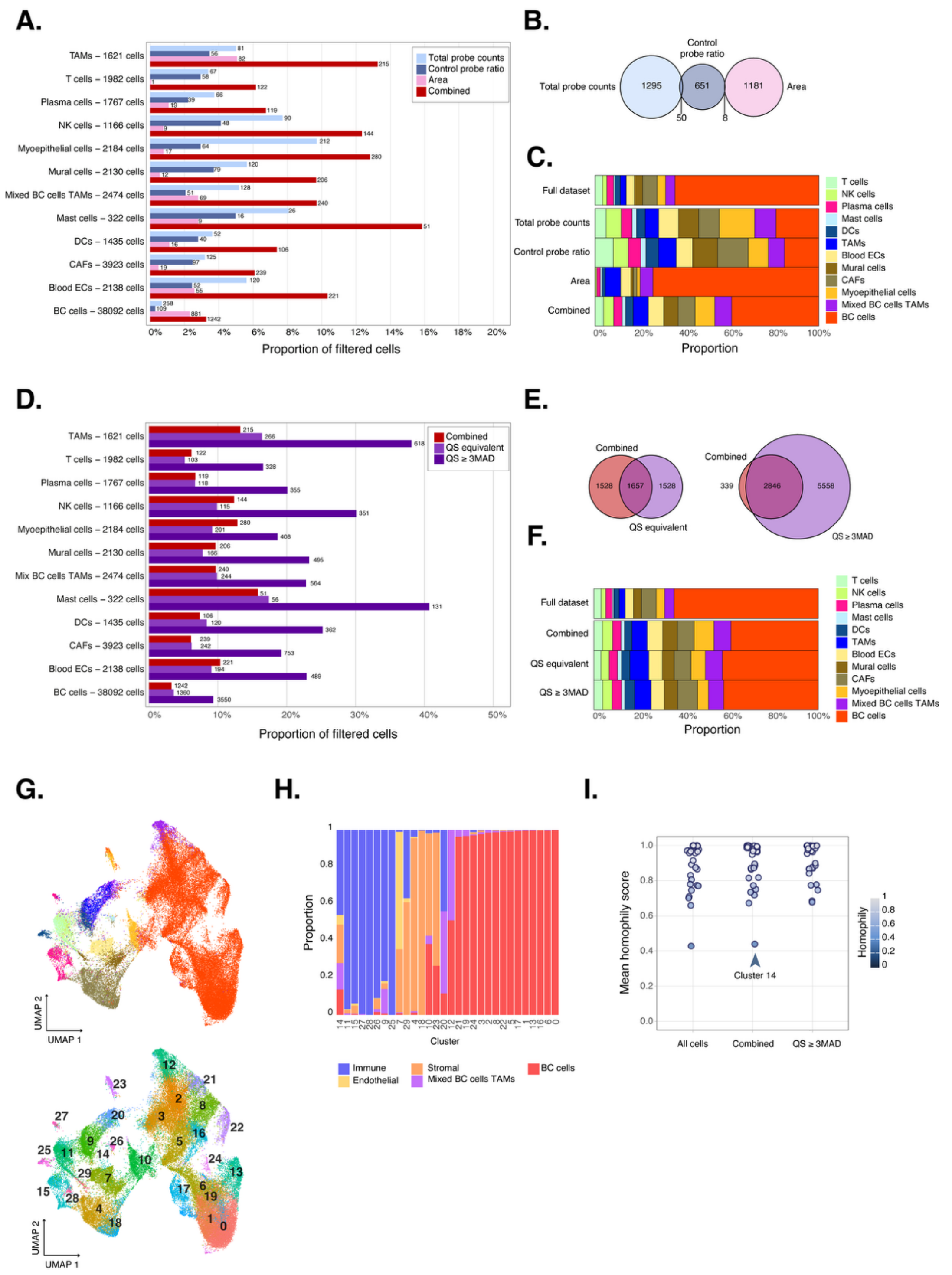

**Supplementary Figure 13. Comparison of SpaceTrooper QS-based filtering with fixed-threshold quality control approaches. A.** Proportion of cells filtered per cell-type using three different quality control criteria, i.e., *total probe counts*, *control probe ratio*, and cell *area*, as well as their combination (*Combined*), following the NanoString AtoMx-recommended workflow. For each cell type and filtering criterion, the number of filtered cells is reported next to the corresponding bar. **B.** Venn diagram showing the overlap between cells filtered by *total probe counts*, *control probe ratio*, and *area*. Overlap between individual criteria is minimal, indicating that each metric captures largely distinct subsets of low-quality cells. No overlap between *total probe counts* and *area* is observed. **C.** Cell-type composition of filtered cells for each individual criterion and for the *combined* approach. The cell-type composition of the full dataset is shown at the top. **D.** Proportion of cells filtered per cell type using the *combined* approach, the *QS equivalent* threshold, and the *QS ≥ 3MAD* threshold. For each cell type and filtering criterion, the number of filtered cells is reported next to the corresponding bar. **E.** Overlap between cells filtered by the *combined* and each QS-based strategy. The *combined* and *QS equivalent* filters show partial overlap, whereas *QS ≥ 3MAD* captures most cells removed by the *combined* approach and identifies additional low-quality cells. **F.** Cell-type composition of cells filtered by the *combined*, *QS equivalent*, and *QS ≥ 3MAD* filtering approaches. The cell-type composition of the full dataset is shown at the top. **G.** UMAP embedding obtained after applying the *combined* filter, colored by cell-type (top) and by cluster identity (bottom). Cluster 14, dispersed across the embedding, still comprises a mixture of all cell lineages. **H.** Proportion of major cell lineages per cluster after applying the *combined* filtering. **I.** Mean homophily score per cluster in all cells and after *combined* or *QS ≥ 3MAD* filtering. After applying the *combined* filter, a cluster with low homophily persists, comparable to that observed in the unfiltered dataset, indicating that a fixed-threshold-based combined approach is insufficient to fully resolve artifactual mixing.

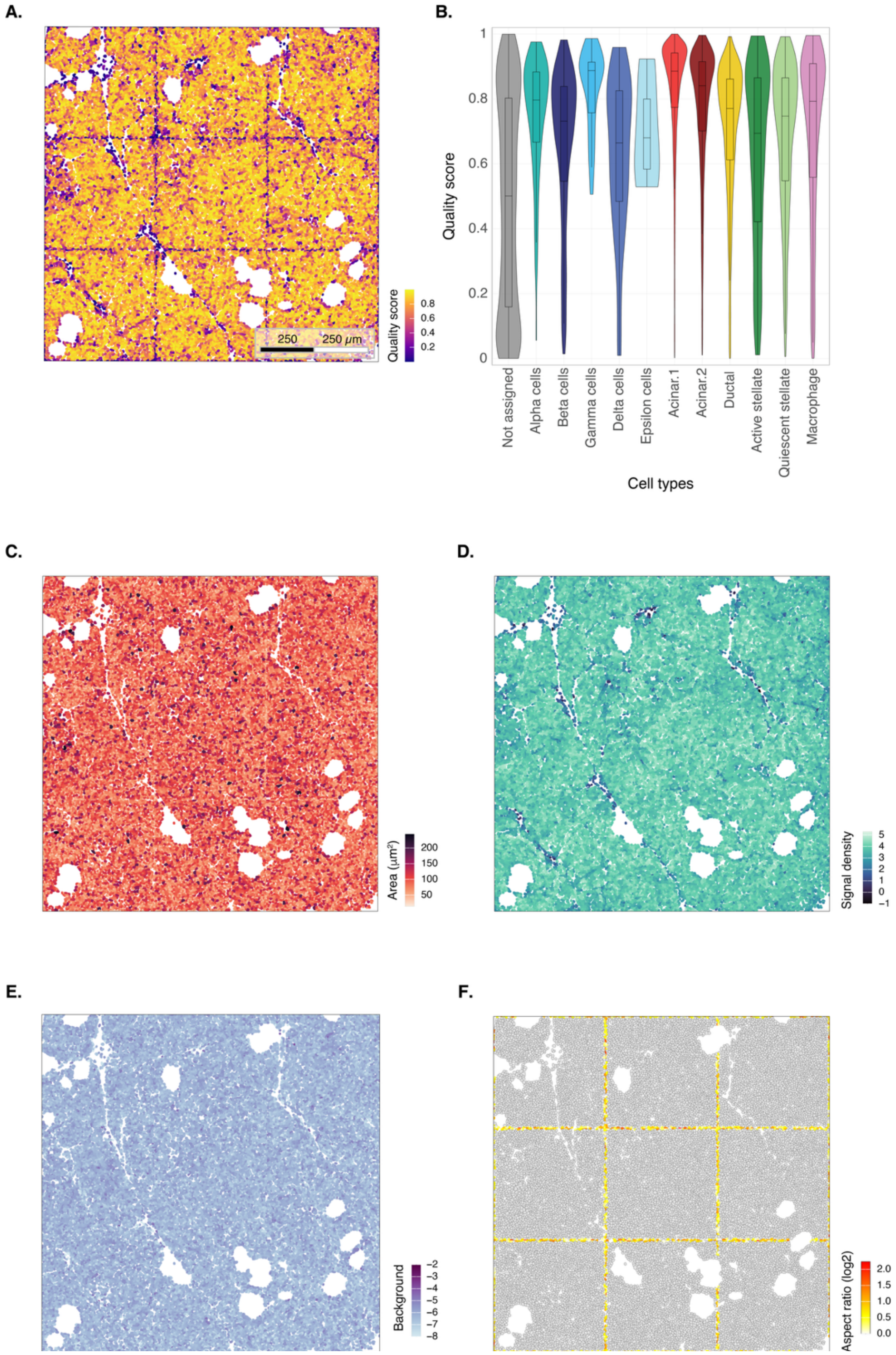


**Supplementary Figure 14. Spatial distribution of QS and QS components in FOVs 51-59 of the CosMx human pancreas dataset. A.** Spatial map of the quality score across the CosMx human pancreas dataset. **B.** Quality score values in the different cell types. Cells labeled as “Not assigned” have been excluded by Nanostring from InSituType phenotyping. **C.** Cell area (in $\mu m^2$). **D.** Signal density. **E.** Background signal. **F.** Absolute $\log_2$ aspect ratio for cells whose centroid lies within 50 px of the nearest FOV boundary.

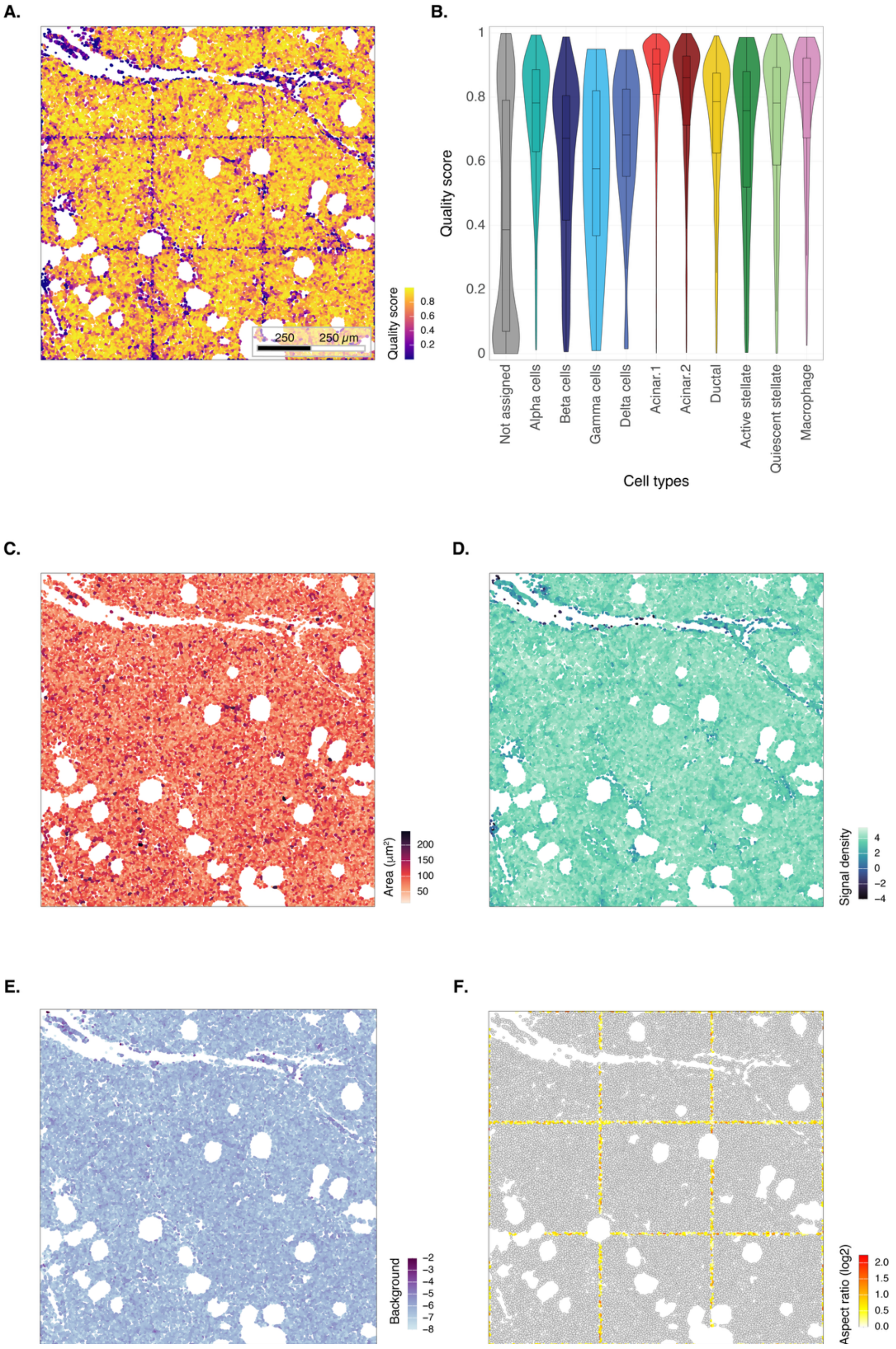


**Supplementary Figure 15. Spatial distribution of QS and QS components in FOVs 60-68 of the CosMx human pancreas dataset. A.** Spatial map of the quality score across the CosMx human pancreas dataset. **B.** Quality score values in the different cell types. Cells labeled as “Not assigned” have been excluded by Nanostring from InSituType phenotyping. **C.** Cell area (in $\mu m^2$). **D.** Signal density. **E.** Background signal. **F.** Absolute $\log_2$ aspect ratio for cells whose centroid lies within 50 px of the nearest FOV boundary.

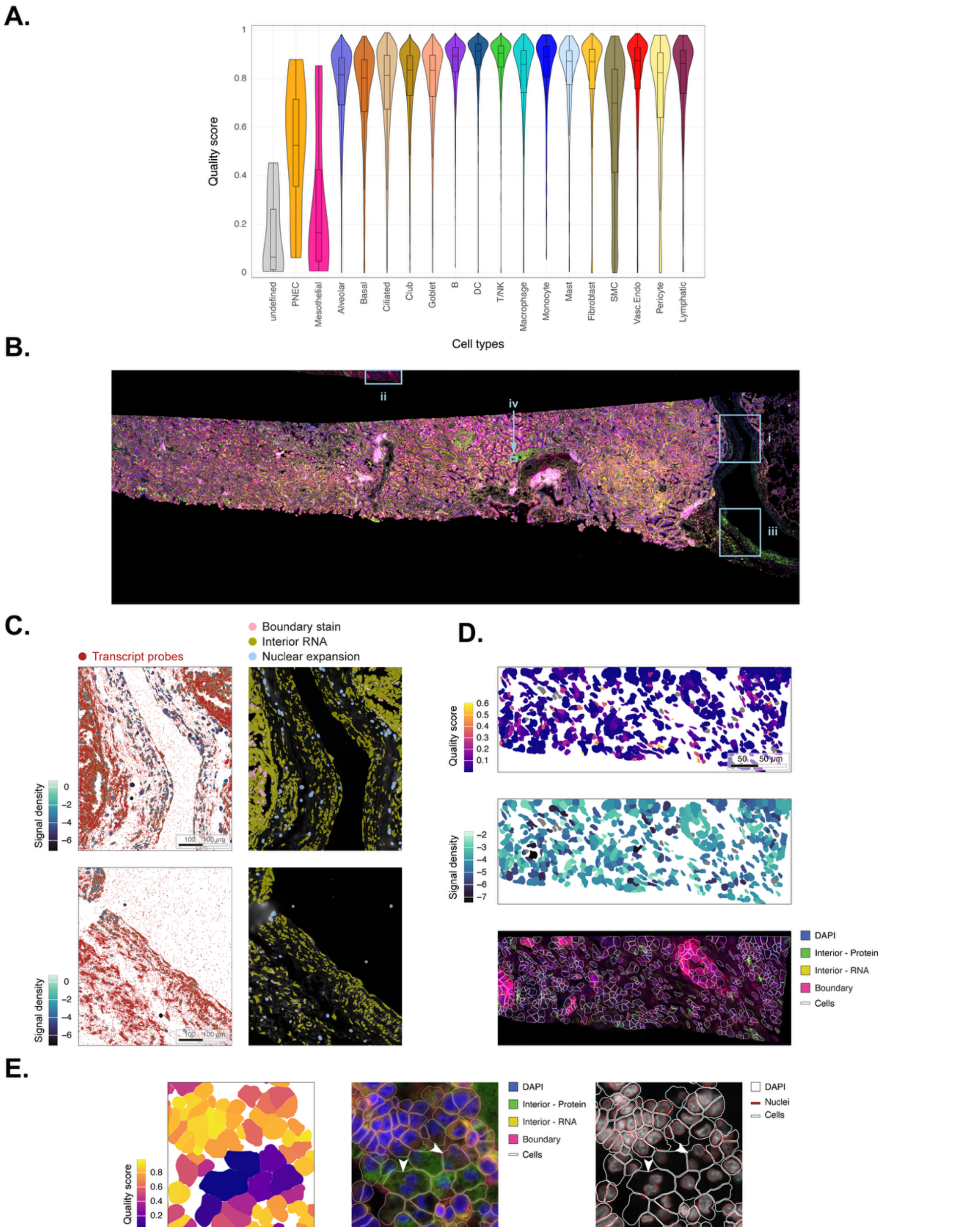


**Supplementary Figure 16. SpaceTrooper QS in the Xenium human lung cancer dataset**. **A.** Quality score values in the different cell types of the Xenium human lung cancer dataset. The *undefined* type comprises 9 cells discarded by the InSituType phenotyping due to very low counts (i.e., 1 and 2). **B.** Composite fluorescence overview of the full tissue section, with boxed regions highlighting areas of interest. **C.** Representative magnifications of two regions in the peripheral vessel (top, boxed region i; bottom, boxed region iii), showing transcript probe localization overlaid on signal density (left) and cells colored by segmentation method (right; boundary stain: pink, interior RNA: dark yellow, nuclear expansion: light blue). **D.** Spatial distributions of the quality score and signal density and composite image of marker staining in the detached tissue patch at the top of the slide (boxed region ii). **E.** Quality score and composite and DAPI images with segmentation overlays in the area identified in **B** as boxed region iv. White arrows indicate spatial multiplets. Composite images have been generated using 10x Genomics Xenium Explorer 4.1.0.

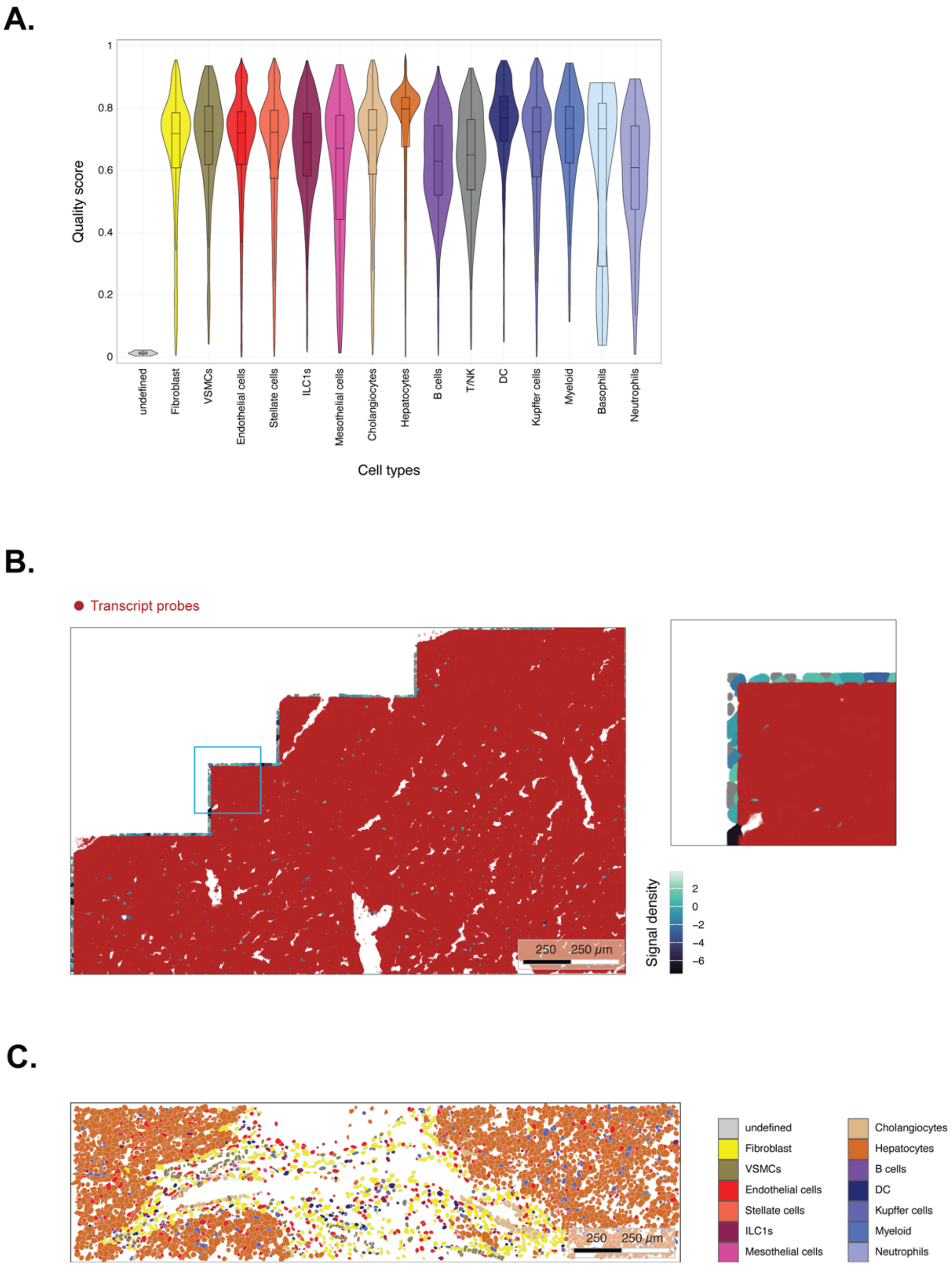


**Supplementary Figure 17. SpaceTrooper QS in the MERFISH mouse liver dataset**. **A.** Quality score values in the different cell types of the Vizgen MERFISH mouse liver dataset. The *undefined* type comprises 5 cells discarded by the InSituType phenotyping due to very low counts (i.e., 1). The sole cell labeled as HsPC has been removed from the violin representation. **B.** Spatial mapping of signal density in a peripheral low-QS region reveals a near-complete depletion of transcript probes (in dark red) along the image border. **C.** Spatial map of cell-type annotations in an interior low-QS region. Cells with zero counts are shown in white.

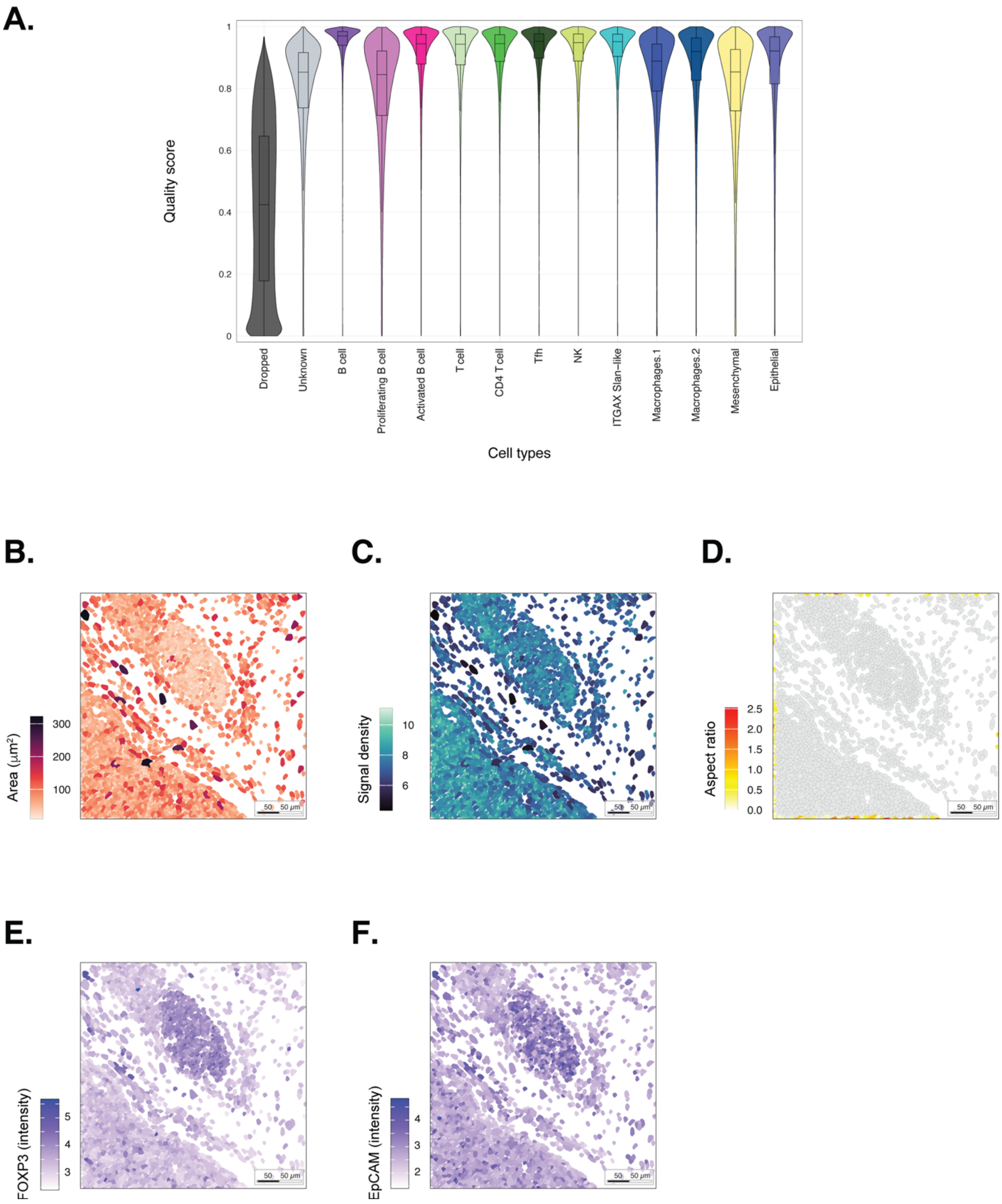


**Supplementary Figure 18. The quality score is generalizable across molecular modalities and captures protein-specific artifacts. A.** Distribution of SpaceTrooper QS across cell types in the CosMx human tonsil protein dataset. **B.** Spatial map of cell area in the region of **Figure 4B**. **C.** Signal density in the same region of **B**. **D.** Absolute $\log_2$ aspect ratio in the same region of **B**. **E.** Spatial distribution of FOXP3 $\log_2$-transformed intensity in the same region of **B**. **F.** Spatial distribution of EpCAM $log_2$-transformed intensity in the same region of **B**.

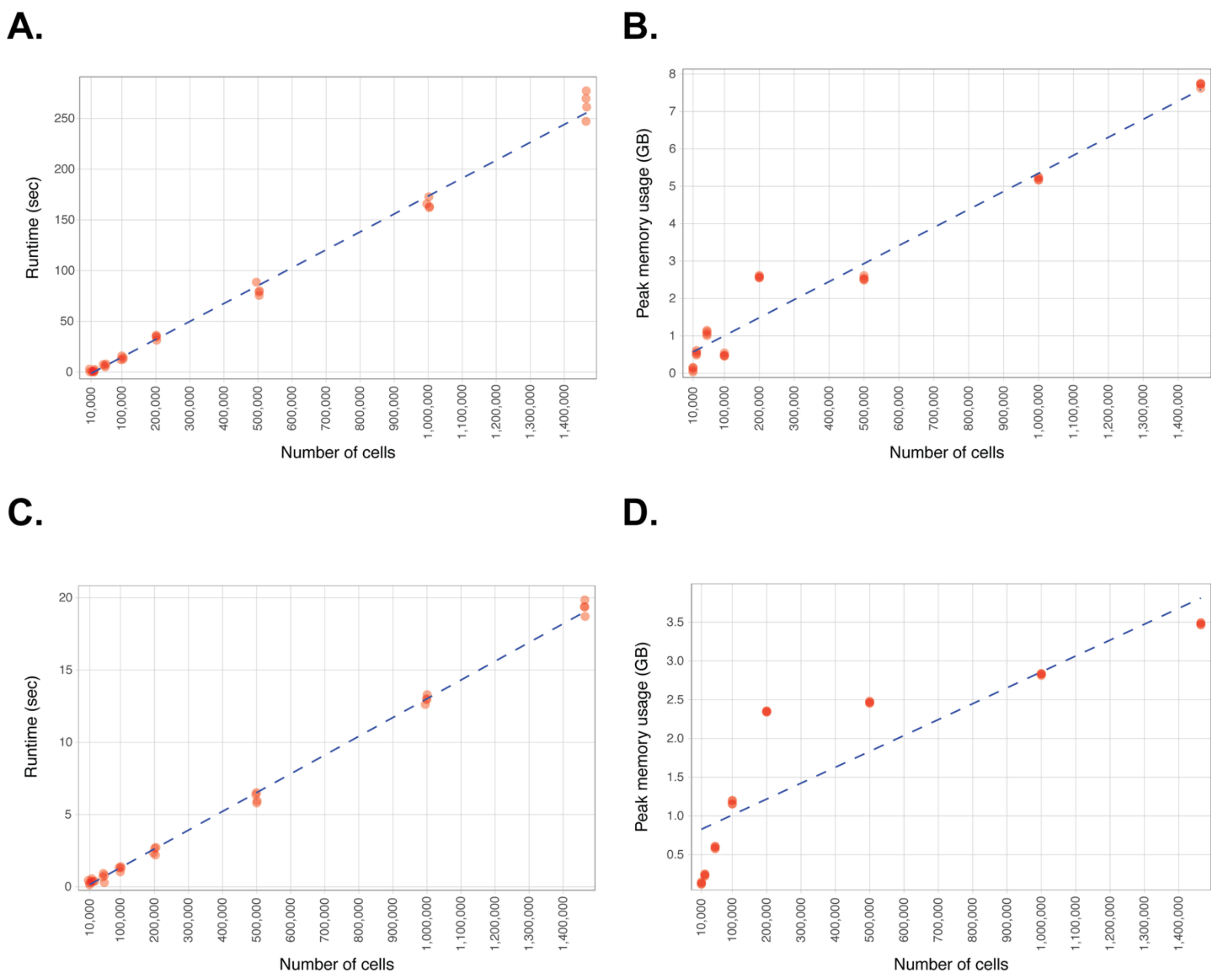


**Supplementary Figure 19. Computing resources and scalability of the entire SpaceTrooper QC frameworks. A.** Runtime of the entire SpaceTrooper QC framework across datasets of increasing size (10,000, 20,000, 50,000, 100,000, 200,000, 500,000, and 1,464,610 cells) generated by random subsampling (n = 4 per size) the CosMx human tonsil protein dataset. The framework runtime is linearly scalable to large datasets, taking less than 300 seconds to process the entire human tonsil dataset. **B.** Peak memory usage of the entire SpaceTrooper QC framework across the same datasets used in panel **A**. **C.** and **D.** same as in **A** and **B** for the QS quantification step alone.
