## Supplementary Tables and Note for "*SpaceTrooper*: a quality control framework for imaging-based spatial omics data"

**Supplementary Table 1:** Cell metadata and metrics available across the imaging-based spatial omics datasets from NanoString CosMx, 10x Genomics Xenium, and Vizgen MERFISH analyzed in this study (✓ available; ✗ not available*).

| Type | Name | Description | CosMx (RNA) | CosMx (protein) | Xenium | MERFISH |
|---|---|---|---|---|---|---|
| Cell and nucleus morphologies | Cell area | Cell area (in pixels assigned to a given cell or in μm$^2$) | ✓ | ✓ | ✓ | ✗ |
| | Nucleus area | Nucleus area (in pixels assigned to a given nucleus or in μm$^2$) | ✗ | ✓ | ✓ | ✗ |
| | Cell width | Cell maximum length along the x dimension (pixels or μm) | ✓ | ✓ | ✗ | ✗ |
| | Cell height | Cell maximum length along the y dimension (pixels or μm) | ✓ | ✓ | ✗ | ✗ |
| | Cell aspect ratio | Cell width divided by cell height | ✓ | ✓ | ✗ | ✗ |
| | Nucleus aspect ratio | Nucleus width divided by nucleus height | ✗ | ✓ | ✗ | ✗ |
| | Cell volume | Cell approximate volume (in μm$^3$) | ✗ | ✗ | ✗ | ✓ |
| | Cell perimeter | Cell perimeter (in pixels) | ✗ | ✓ | ✗ | ✗ |
| | Circularity | Cell area divided by cell perimeter (1: circular; <1: less circular) | ✗ | ✓ | ✗ | ✗ |
| | Eccentricity | Cell minor axis divided by cell major axis | ✗ | ✓ | ✗ | ✗ |
| | Solidity | Cell area divided by cell convex area (a measure of cell regularity with values <1 indicating increased cell irregularity) | ✗ | ✓ | ✗ | ✗ |
| Molecular content | Total probe count | Total probe count per cell (RNA) or total intensity per cell (protein) | ✓ | ✓ | ✓ | ✓ |
| | Total uniquely detected probes | Total number of uniquely detected probes per cell | ✓ | ✓ | ✓ | ✓ |
| Background noise | Total negative control probe count | Total negative control probe counts per cell | ✓ | ✓ | ✓ | ✓ |
| | Total uniquely detected negative control probes | Total number of uniquely detected negative control probes per cell | ✓ | ✓ | ✓ | ✓ |
| | Total control code words count | Total control code word counts per cell | ✗ | ✗ | ✓ | ✗ |
| | Total uniquely detected control code words | Total number of uniquely detected control code words per cell | ✗ | ✗ | ✓ | ✗ |
| | Total negative control code words count | Total negative control code word counts per cell | ✓ | ✗ | ✓ | ✗ |
| | Total uniquely detected negative control code words | Total number of uniquely detected negative control code words per cell | ✓ | ✗ | ✓ | ✗ |
| FOV-related | Cell local position | Cell position within the FOV | ✓ | ✓ | ✗ | ✗ |
| | Split ratio | The ratio of cell area to mean cell area in each FOV if the cell touches the FOV border, 0 otherwise | ✗ | ✓ | ✗ | ✗ |
| FOV-level statistics | Median target probe expression | Median RNA target probe expression across all cells within a given FOV | ✗ | ✓ | ✗ | ✗ |
| | Percentile target probe expression | Percentiles (e.g., 75, 80, 85, etc.) of RNA target probe expression across all cells within a given FOV | ✗ | ✓ | ✗ | ✗ |
| | Median negative probe expression | Median negative probe expression across all cells within a given FOV | ✗ | ✓ | ✗ | ✗ |
| | Percentile negative probe expression | Percentiles (e.g., 75, 80, 85, etc.) of negative probe expression across all cells within a given FOV | ✗ | ✓ | ✗ | ✗ |
| | Fraction of unassigned transcripts | Proportion of transcripts in a FOV that are not assigned within any cell of the FOV | ✓ | ✗ | ✗ | ✗ |
| Fluorescence signals | Mean fluorescence intensity of a morphological marker | Mean fluorescence intensity of a given marker in each cell | ✓ | ✓ | ✗ | ✗ |
| | Maximum fluorescence intensity of a morphological marker | Maximum fluorescence intensity of a given marker in each cell | ✓ | ✓ | ✗ | ✗ |

*Although not provided in the metadata of the analyzed datasets, some of the cell and nucleus morphologies metrics can be computed from the cell segmentation polygons.

**Supplementary Table 2:** Overview of dataset summary metrics (including total cells, total probes, cells with zero total counts, and cells lacking control probe counts), methods and thresholds used to identify high- and low-quality examples for each predictor component, and GLM model-fitting parameters used to quantify the quality score (QS) across the spatial omics datasets examined in this study.

| | | | CosMx (RNA) *Human breast cancer* | CosMx (RNA) *Human pancreas* | Xenium | MERFISH | CosMx (protein) *Human tonsil* |
|---|---|---|---|---|---|---|---|
| **General summary** | Total cells | | 59,284 | 48,944 | 162,254 | 395,215 | 1,464,466 |
| | Total probes | | 1,010 | 21,731 | 541 | 385 | 69 |
| | Cells with total count equal to 0 | | 50 | 0 | 360 | 1332 | 0 |
| | Cells with control probes counts equal to 0 | | 43,366 | 231 | 161,014 | 92,923 | 0 |
| **Cell size** | Outlier detection method | | Medcouple | 3MAD | Medcouple | Medcouple | Medcouple |
| | Threshold high | | 252.94 $\mu m^2$ | 168.13 $\mu m^2$ | 283.30 $\mu m^2$ | 3,739.02 $\mu m^3$ | 162.07 $\mu m^2$ |
| | Low-quality examples | | 400 | 358 | 437 | 3* | 18,143 |
| **Signal density** | Outlier detection method | | Medcouple | Medcouple | Medcouple | 1st percentile | 3MAD |
| | Threshold low | | -3.10 | 2.24 | -2.61 | -4.97 | 13.47 |
| | Low-quality examples | | 337 | 885 | 3,147 | 3,939 | 36,594 |
| **Background signal** | Outlier detection method | | Medcouple | 3MAD | 3MAD | 3MAD | 3MAD |
| | Threshold high | | -5.13 | -4.86 | -2.62 | -5.45 | -8.64 |
| | Low-quality examples | | 184 | 195 | 13* | 3585 | 28289 |
| **Border effect** | Outlier detection method | | 3MAD | 3MAD | --- | --- | 3MAD |
| | Thresholds (low÷high) | | -1.168÷1.197 | -1.086÷1.057 | --- | --- | -1.126÷1.126 |
| | Low-quality examples (low÷high) | | 206÷223 | 177÷207 | --- | --- | 4,306÷3,947 |
| **Model fitting** | Total low-quality examples | | 1,333 | 1,733 | 3,566 | 7,419 | 72,993 |
| | Total high-quality examples | | 1,333 | 1,733 | 3,566 | 7,419 | 72,993 |
| | Intercept | | -1.07 | -3.54 | 2.98 | -0.56 | -28.04 |
| | $\beta_i$ | Cell size | -0.01 | -0.01 | -0.01 | NA* | -0.01 |
| | | Signal density | 0.55 | 1.01 | 1.08 | 0.56 | 1.04 |
| | | Background signal | -0.29 | 0.03 | NA* | -0.21 | -0.99 |
| | | Border effect | -1.59 | -1.17 | --- | --- | -1.29 |
| | $\beta_{ij}$ | Cell size:Signal density | 0 | 0 | 0 | NA | 0 |
| | | Cell size:Background signal | 0 | 0 | NA | NA | 0 |
| | | Cell size:Border effect | -0.01 | 0 | --- | --- | -0.01 |
| | | Signal density:Background signal | -0.04 | -0.14 | NA | -0.03 | -0.05 |
| | | Signal density:Border effect | -0.45 | -0.35 | --- | --- | -0.09 |
| | | Background signal:Border effect | 0.13 | 0.18 | --- | --- | 0.14 |

*The predictor component was excluded from the QS calculation because fewer than 0.1% of all cells were identified as low-quality examples by this metric.

### Supplementary Note 1: Selection of the quality score components

Quality control (QC) and data filtering of imaging-based spatial omics data are generally guided by criteria recommended by the respective technology manufacturers[1-4]. For NanoString CosMx, it is recommended to exclude cells whose area represents an extreme outlier, as well as cells with low transcript abundance (typically fewer than 20 detected transcripts for 1000-plex assays or fewer than 50 for 6000-plex assays) and cells with a high proportion of negative control counts (>1% of total counts per cell), which collectively reflect segmentation artifacts and insufficient molecular signal (see

NanoString online documentation for details; https://nanostring-biostats.github.io/CosMx-Analysis-Scratch-Space/ and https://university.nanostring.com/cosmx-smi-data-analysis-user-manual). For 10x Genomics Xenium and Vizgen MERFISH, no strictly universal QC thresholds are defined, as optimal cutoffs strongly depend on tissue type, gene panel design, and sample quality. For Xenium, 10x Genomics explicitly acknowledges the absence of universal thresholds but recommends filtering cells with very low transcript counts (typically <10 total transcripts and <5 unique transcripts), elevated negative control probe fractions (>5% per control probe set per cell), or low decoded nuclear transcript density (<1 transcript per 100 $\mu m^2$). In MERFISH experiments, cells with extreme sizes (e.g., volumes <200 $\mu m^3$ or >3,000 $\mu m^3$) or with very low transcript counts (commonly <10-50 transcripts per cell, depending on the panel) are frequently removed.
Although these manufacturer-guided QC metrics effectively mitigate technical artifacts and promote standardized analyses across platforms, several of them are intrinsically interdependent. To systematically assess standard QC metrics and their relationships, we examined the cell-level metadata provided by NanoString CosMx, 10x Genomics Xenium, and Vizgen MERFISH in a set of publicly available datasets processed using the manufacturers' standard pipelines. These metadata include morphological features of cells and nuclei, measurements of target and control probe signals, and spatial attributes relative to the field of view (FOV) (**Supplementary Table 1**). From the metrics shared across all three technologies, we focused on cell size (cell area or cell volume, as for MERFISH), signal density (defined for RNA assays as total cell counts normalized by cell size and for protein assays as the sum of average protein intensities) and background signal (quantified as the ratio of control counts to total counts). In addition, because standard CosMx pipelines process each FOV independently, we further analyzed the cell aspect ratio (width-to-height) in relation to the distance of the cell centroid from the nearest FOV border to capture potential border-related artifacts.

*Cell size (area or volume)*. Cell size (area or volume) is a critical parameter for interpreting transcript counts at the single-cell level. In CosMx, Xenium, and MERFISH, cell boundaries are defined using segmentation algorithms based on the staining of morphological markers. Although these technologies differ in imaging chemistry and segmentation strategies, the distribution of cell sizes within a given sample is largely determined by tissue composition and cellular heterogeneity, as different cell types exhibit characteristic dimensions and morphologies. Consistent with recent reports[2,5], we observed that cell sizes across all analyzed samples are predominantly below 200-300 $\mu m^2$ for cell area and 2000-2500 $\mu m^3$ for cell volume, with median values lying between 50-100 $\mu m^2$ and approximately around 600 $\mu m^3$ for area and volume, respectively (**Supplementary Figures 1A, 1C, 1E, and 4A**). Across all samples and tissue types, the cell-size distribution displays a right-skewed tail extending toward very large values. While cells with very small areas may arise from over-segmentation, they often correspond to genuinely small cell types (e.g., some immune cells). In contrast, although large cells are biologically plausible, such as muscle cells or mature adipocytes, they are rarely detected in imaging-based spatial omics datasets, likely because segmentation algorithms often fail to delineate their irregular morphologies or their cytoplasm-poor anatomy in thin tissue sections[6-10]. Moreover, a single large polygon may occasionally encompass multiple adjacent cells or include both a cell and portions of its surrounding environment, while cell debris and fluorescence artifacts may also be misidentified as cells. In all such cases, expression quantification becomes unreliable and may not accurately reflect a true cellular identity. This interpretation is further supported by the relationship between cell size and the total number of measured mRNA molecules. Within the biologically expected range of cell sizes, we observe an approximately linear correlation between these two metrics; however, beyond this range, the correlation is no longer consistently preserved across all RNA assays (**Supplementary Figures 1B, 1D, and 1F**) and in the CosMx protein assay, where the total signal is quantified as summed protein intensity rather than transcript counts (**Supplementary Figure 4B**). Consequently, to capture segmentation-related artifacts characterized by implausibly large cells while preserving genuinely small cell types, we identified cell-size outliers in the right tail of the cell-size distribution.

*Signal density*. Measurements of total transcript counts per cell are influenced by many of the same factors that affect cell size, including cell-type specific expression programs, tissue composition, and segmentation artifacts. In addition, total counts also depend on the size of the targeted gene panel, which in the analyzed datasets ranges from a few hundred genes to whole-transcriptome panels. Inspection of the total probe count distributions reveals substantial variability across samples, with distinct median values and the consistent presence of cells with very low transcript numbers (**Supplementary Figures 2A, 2C, and 2E**). As is commonly done in droplet-based single-cell RNA-seq analyses, one could apply a hard threshold on total counts to discard low-quality cells. However, overly stringent cutoffs risk removing small, but biologically valid, cells with limited yet informative transcriptome content, rather than selectively excluding empty or dead cells. To overcome this limitation, we leveraged the concurrent availability of cell morphology information and quantified the signal density, defined as the total number of counts normalized by cell size (**Supplementary Figures 2B, 2D, and 2F**). Thresholding on the density distribution, rather than on total counts alone, enables the selective exclusion of cells with abnormally low molecular signal while preserving genuinely small cells. This is particularly relevant because transcript abundance may reflect not only cell identity but also the specific 2D section of the cell captured in the tissue. For the CosMx protein dataset (**Supplementary Figure 4C**), the signal density, quantified as the summed average protein intensity, exhibits a higher overall signal with low-signal outliers evident in the left tail of the distribution.

*Background signal*. Control probes differ in number and design across technologies: they may be physical probes that do not match any genomic sequence or codewords with no corresponding physical probe. Because these probes capture background levels and non-specific signal in the decoding process, they provide a measure of background noise. To quantify the background signal, we combined negative control probes and blank controls depending on their availability across technologies[2]. In RNA assays, a substantial fraction of cells exhibits zero control probe counts, with this proportion varying across technologies (**Supplementary Table 2**). Focusing on cells with non-zero control probe counts, we calculated the proportion of control probe counts to total counts per cell. This proportion is generally low but exhibits a right-skewed distribution in all samples (**Supplementary Figures 3A, 3C, and 3E**). Cells with high proportions of control probe counts are mostly those with low total counts, though some exceptions exist (**Supplementary Figures 3B, 3D, and 3F**). Irrespective of total counts, cells with elevated background fractions have expression profiles substantially influenced by non-specific signals and may therefore be less reliable for downstream analyses. In the

CosMx protein assay, two non-specific antibodies (mouse IgG1 and rabbit IgG) are used as control probes. Because the assay measures averaged fluorescence per cell, even faint signals are detected, and no cells exhibit zero control probe intensity (**Supplementary Table 2**). Nonetheless, the distribution of the proportion of control probe intensity remains right-skewed (**Supplementary Figure 4D**), and scatterplots show the expected inverse relationship: cells with higher background proportions generally have lower total signal intensity (**Supplementary Figure 4E**).

*Border effect*. The field of view (FOV) corresponds to the tissue region captured by the imaging system, and FOVs are processed differently across spatial omic technologies. In particular, the standard NanoString CosMx pipeline processes each FOV independently, which can lead to artifacts such as fractured or distorted cells near FOV borders. These artifacts may affect segmentation accuracy and, consequently, gene or protein quantification. Consistent with this, we observed that both cell area and total probe counts (or total intensity) tend to decrease as cells approach the nearest FOV border (**Supplementary Figures 5A, 5B, 6A, and 6B**). To selectively identify potentially distorted cells without discarding all border-proximal cells, we leveraged the cell aspect ratio, defined as the ratio of cell width to height (**Supplementary Table 1**). For approximately circular cells, this metric is expected to be close to 1 ($\log_2$ aspect ratio equal to 0). Plotting $\log_2$ aspect ratio against the distance from horizontal or vertical borders shows that cells maintain values near 0 at larger distances but deviate significantly when close to FOV edges. Specifically, the aspect ratio increases near horizontal borders due to reduced cell height (**Supplementary Figures 5C and 6C**) and decreases near vertical borders due to reduced cell width (**Supplementary Figures 5D and 6D**). In the analyzed datasets, these deviations begin roughly in an interval of 40-80 pixels from the FOV border. Below this threshold, we observe outliers with extreme aspect ratios. These observations support the use of the aspect ratio as a robust metric to identify cells potentially distorted by the border effect.

Across all QC metrics (i.e., cell size, signal density, background signal, and border effect), outliers can be identified either by applying predefined thresholds or by examining the empirical distributions of the data. Analysis of these distributions across the selected samples highlights two key considerations: first, outlier thresholds should be adaptive and reflect dataset-specific variability (for example, using median absolute deviation or Medcouple-adjusted statistics; see **Supplementary Figures 1-6**) rather than fixed absolute cutoffs; second, the effectiveness of outlier detection can vary across samples, underscoring the importance of combining multiple complementary approaches and evaluating their performance on a per-dataset basis.

### REFERENCES

1 Marco Salas, S. *et al.* Optimizing Xenium In Situ data utility by quality assessment and best-practice analysis workflows. *Nat Methods* **22**, 813-823 (2025). https://doi.org:10.1038/s41592-025-02617-2

2 Ozirmak Lermi, N. *et al.* Comparison of imaging based single-cell resolution spatial transcriptomics profiling platforms using formalin-fixed paraffin-embedded tumor samples. *Nat Commun* **16**, 8499 (2025). https://doi.org:10.1038/s41467-025-63414-1

3 Martin, N. *et al.* MerQuaCo: a computational tool for quality control in image-based spatial transcriptomics. (2025). https://doi.org:10.7554/elife.105149.1
4 Pitino, E. *et al.* STAMP: Single-cell transcriptomics analysis and multimodal profiling through imaging. *Cell* **188**, 5100-5117 e5126 (2025). https://doi.org:10.1016/j.cell.2025.05.027
5 Liu, J. *et al.* Concordance of MERFISH spatial transcriptomics with bulk and single-cell RNA sequencing. *Life Sci Alliance* **6** (2023). https://doi.org:10.26508/lsa.202201701
6 Wang, H. *et al.* Systematic benchmarking of imaging spatial transcriptomics platforms in FFPE tissues. *Nat Commun* **16**, 10215 (2025). https://doi.org:10.1038/s41467-025-64990-y
7 Petukhov, V. *et al.* Cell segmentation in imaging-based spatial transcriptomics. *Nat Biotechnol* **40**, 345-354 (2022). https://doi.org:10.1038/s41587-021-01044-w
8 Bruhns, M. *et al.* Effects of segmentation errors on downstream-analysis in highly-multiplexed tissue imaging. *PLoS Comput Biol* **21**, e1013350 (2025). https://doi.org:10.1371/journal.pcbi.1013350
9 Mitchel, J., Gao, T., Cole, E., Petukhov, V. & Kharchenko, P. V. Impact of Segmentation Errors in Analysis of Spatial Transcriptomics Data. *bioRxiv*, 2025.2001.2002.631135 (2025). https://doi.org:10.1101/2025.01.02.631135
10 Pang, M., Roy, T. K., Wu, X. & Tan, K. CelloType: a unified model for segmentation and classification of tissue images. *Nat Methods* **22**, 348-357 (2025). https://doi.org:10.1038/s41592-024-02513-1
